# Spatial subdomains in the Optic Tectum for the encoding of visual information

**DOI:** 10.1101/2023.05.15.540762

**Authors:** Thomas Shallcross, Giovanni Diana, Juan Burrone, Martin Meyer

## Abstract

Neurons across the visual system provide estimates of the visual features they encode. However, the reliability of those estimates can vary across the neuronal population. Here, we use information theory to provide a spatial map of how well neurons can distinguish ethologically-relevant visual stimuli across the entire larval zebrafish optic tectum, a brain region responsible for driving visually guided behaviour. We find that the ability of neurons to discriminate between stimuli is non-uniformly distributed across the tectum. Specifically, we show that information about local motion is preferentially encoded in the posterior tectum, whilst information about whole-field motion is preferentially encoded in the anterior tectum. This is achieved through two systematic changes along the anterior-posterior axis of the tectum: (i) a change in the number of neurons that discriminate between stimuli and (ii) a change in how well each neuron can discriminate between stimuli. By classifying neurons into distinct subtypes based on their response properties we uncovered a small group of neurons that are spatially localised to specific regions of the tectum and are able to discriminate between visual stimuli in a highly reliable manner. Our results highlight the importance of implementing information theoretic approaches to assess visual responses and provide a novel description of regional specialisation in the zebrafish optic tectum.

## 2 Introduction

Our perception of the visual world is shaped by the information which is transmitted from our eyes to the rest of the brain via the axons of the retinal ganglion cells (RGCs). A major target of the RGCs is the optic tectum/superior colliculus which is involved in guiding movements to specific points in body-centred space (Basso and May 2017, Cooper and Mcpeek 2021, Isa et al. 2021). Across all species investigated there is an ordered retinotopic mapping of connections from the eye to the tectum, such that adjacent regions of visual space are represented by adjacent tectal neurons (Cang and Feldheim 2013, Isa et al. 2021). Furthermore, the RGCs split up the representation of visual space into multiple segregated channels of information, with each channel dedicated to processing specific visual features, such as direction of motion or object size (Lettvin et al. 1959, Kerschensteiner and Hardesty 2022). This means the tectum receives input from multiple, parallel, retinotopically organised ‘feature’ maps. However, understanding how the tectum processes these features, and how they relate to behaviour is an ongoing area of research (Basso and May 2017, Cooper and Mcpeek 2021).

The larval Zebrafish has served as a particularly useful model organism for understanding how the tectum encodes these visual features due to the animal’s small size, optical transparency, and ability to elicit behaviours using simple visual stimuli (Vanwalleghem et al. 2018). Previous research has demonstrated the presence of a number of feature detectors within the tectum such as direction, size, and orientation selective responses (Bianco and Engert 2015, Del Bene et al. 2010, Hunter et al. 2013), which can either be inherited directly from the RGC inputs or generated *de novo* in the tectum (Förster et al. 2020, Hunter et al. 2013). Initially, it was thought that each feature detector subtype was uniformly distributed throughout the tectum, allowing each visual feature to be uniformly represented across visual space (Del Bene et al. 2010, Niell and Smith 2005, Preuss et al. 2014, Romano et al. 2015, Wang et al. 2019). This was supported by the fact that the spatial distribution of neuronal subtypes within the tectum demonstrated a lack of stereotypy across fish, implying there is no regional specialisation in the tectum (Chen et al. 2018, Portugues et al. 2014). On the other hand, more recent work has shown both size and direction-selective neurons which appear to be spatially localised to specific regions of the tectum, indicating there are retinotopic biases in how visual features are encoded across visual space (Förster et al. 2020). These regional specialisations within the tectum are thought to reflect changes in the local statistics of the animal’s sensory environment (Alexander et al. 2022, Förster et al. 2020, Wang et al. 2020). Moreover, it is thought that the behavioural relevance of visual stimuli can change across visual space (Muto et al. 2013), raising the possibility that different regions of the tectum are adapted to encode different types of behaviorally relevant visual information. Therefore, in the present study, we sought to understand whether there are retinotopic biases in how the tectum encodes information to different types of visual stimuli. In particular, we compare neurons that encode information about local and whole-field motion. Both types of motion have previously been used in separate studies to probe for feature selectivity (Bianco and Engert 2015, Chen et al. 2018). However, at the behavioural level, local and whole-field motion represent fundamentally different types of visual information: local motion stimuli are able to induce approach behaviours (Bianco et al. 2011, Semmelhack et al. 2014), whilst whole-field motion stimuli induce stabilisation reflexes (Brockerhoff et al. 1995, Kubo et al. 2014, Naumann et al. 2016). Since regional specialisations within the tectum are thought to reflect adaptations to behaviorally relevant visual stimuli, we wanted to address whether there were regional differences in how the tectum encodes these two different types of motion.

Previous studies that have looked for regional specialisations across brain regions have typically focused on the non-uniform distribution of neuronal subtypes (Heukamp et al. 2020). However, it is not only the non-uniform distribution of neuronal subtypes that could lead to regional specialisation. Since single neurons are known to be unreliable encoders of information due to trial-to-trial response variability (Mochol et al. 2010, Ruyter van Steveninck et al. 1997, Stringer et al. 2021) it is also possible that the reliability with which neurons respond to specific visual stimuli changes systematically throughout the tectum. This in turn could lead to specific regions of the tectum containing neurons that encode higher amounts of information about certain visual features (Avitan et al. 2020). Therefore, to fully understand whether the tectum contains regional specialisations it is important to answer two questions: 1) what is the spatial distribution of neurons that encode information about visual stimuli? 3) does the reliability with which neurons discriminate between visual stimuli change systematically across the tectum?

To allow us to address the above questions we take an information theoretic approach. Information theory provides a way to quantify the mutual dependence between two variables (Shannon 1948), such as whether a neuron’s response correlates with the presentation of a particular stimulus, and therefore, can be used to quantify how reliably a neuron can discriminate between groups of stimuli. Consistent with a previous study (Förster et al. 2020) we find that the optic tectum does not uniformly encode visual features along its anterior-posterior axis. In particular, we show that the posterior region of the tectum is enriched with neurons that can discriminate between local motion stimuli, whilst the anterior region of the tectum is enriched with neurons that can discriminate between whole-field motion stimuli. We further show that these results can be explained by the spatial localisation of a number of neuronal subtypes each of which can discriminate between certain features of the stimuli. Overall, our results show that changes in how reliably neurons can discriminate between visual stimuli contributes to regional specialisations within the tectum.

## 3 Results

### 3.1 Calculating the information content of neuronal responses to local and whole-field motion

To analyse visually evoked response properties of neurons in the optic tectum we performed 2-photon calcium imaging on transgenic Zebrafish larvae expressing nuclear localised pan-neuronal GCaMP6s (elavl3:H2B-GCaMP6s) at 7 days post fertilisation (dpf) across one tectal hemisphere, whilst showing visual stimuli to the contralateral eye (Figure 1A & B). Cells were segmented and calcium transients were extracted using suite2p (Pachitariu et al. 2017, Figure 1C & D) and then classified as to whether or not they were visually responsive (Figure S1). We found 1043 ± 129 neurons per fish (n = 9 fish) were classified as visually responsive which represents 46% ± 7% of all segmented neurons (Figure 1F). To understand the response properties of these visually responsive neurons we took an information theoretic approach by utilising a metric known as mutual information (MI) which quantifies how well a neuron can discriminate between multiple stimuli. Neurons which have high mutual information can be thought of as having well-separated response distributions to different visual stimuli and are therefore more able to discriminate between the stimuli. Those that have low mutual information have more overlapping response distributions and are less able to discriminate between stimuli (Figure 1E). We were interested in comparing neurons which encoded information about i) local motion stimuli (dots) or ii) whole-field motion stimuli (gratings) (Figure 1B and S3A-C). Each local motion stimulus was matched to a corresponding whole-field motion stimulus such that the features of the stimuli were the same in terms of the direction of motion, speed, and size (the diameters of the dots were matched to the wavelengths of gratings). This ensured we could compare the same feature selectivity across local and whole-field motion. For each neuron, we calculated the mutual information to both local and whole-field motion stimuli. We then found the neurons which had non-zero mutual information to either local motion or whole-field motion. Overall we found 224 ± 49 neurons encoded information about local motion, whilst 171 ± 31 neurons encoded information about whole-field motion (Figure 1G). In this paradigm, the maximum number of bits a neuron could encode is 2 bits. If a neuron were to encode 2 bits of information it would mean that the response distributions to all stimuli were non-overlapping. Neurons encoding less than 2 bits provide only partial information about the stimuli due to overlapping response distributions to the different visual stimuli (Figure S2). The average amount of information encoded was 0.31 ± 0.05 bits vs 0.27 ± 0.02 bits of information to the local motion and whole-field motion stimulus sets, respectively (Figure 1H). In both cases, most neurons encode only a small number of bits, with only a few neurons encoding over 1 bit of information (Figure 1I and J).

**Figure 1:**
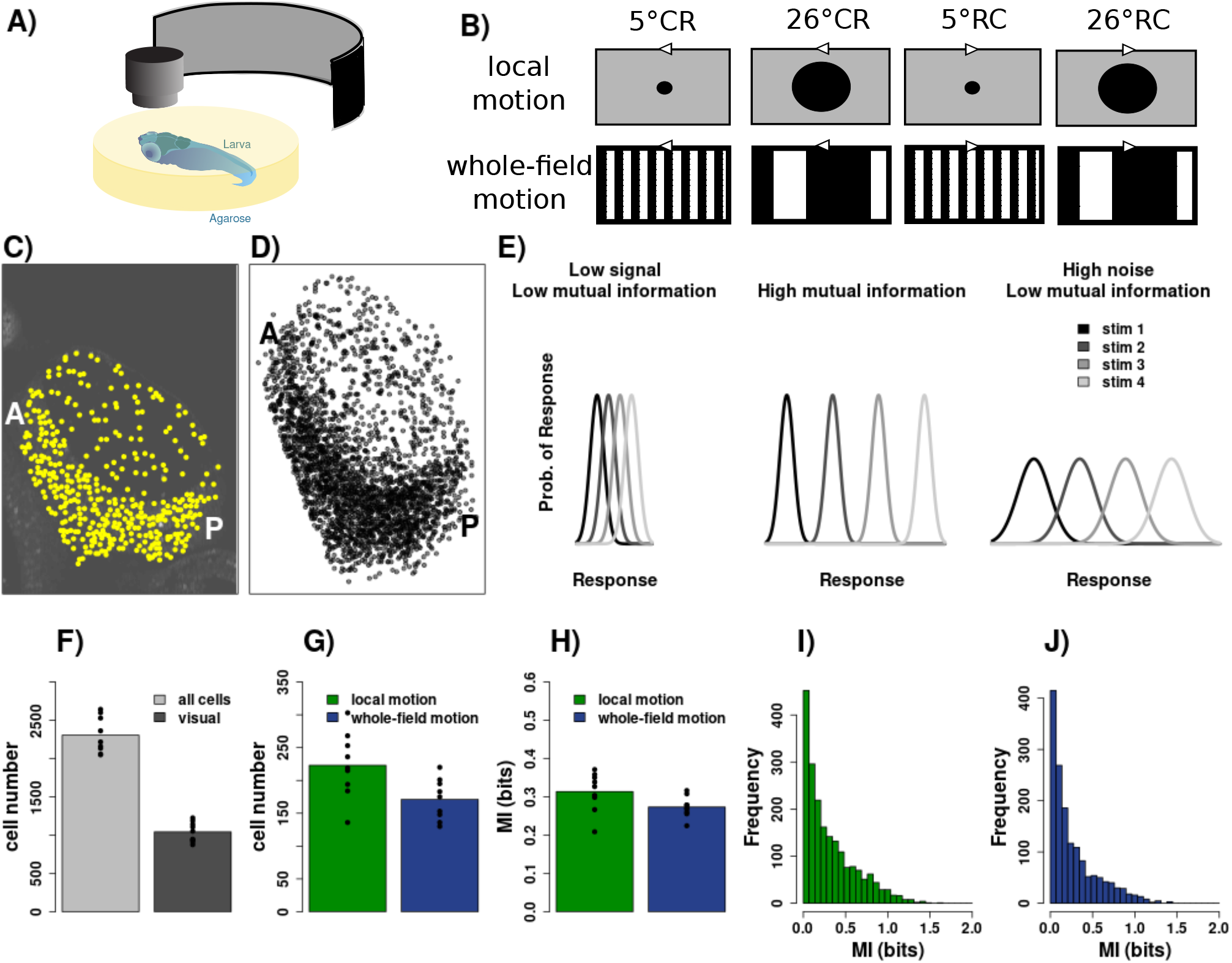
Mutual information of neuronal responses. (**A**) Schematic of functional imaging set up. (**B**) The two sets of visual stimuli -local motion (dots), and whole-field motion (sinusoidal gratings). Both stimulus sets are composed of 4 stimuli which vary in size and direction (5° and 26° of visual angle which corresponds 1/5 cycles/° and 1/26 cycles/° spatial frequency for the gratings. CR: caudal to rostral motion, RC: rostral to caudal motion). (**C**) An example plane of the tectum overlaid with the location of the segmented neurons (A = anterior, P = posterior). (**D**) Segmented neurons from all 7 imaging planes in one example fish. (**E**) The information content of neurons depends on both the magnitude of the response to each stimulus (signal) as well as the variability of the response to the stimuli (noise). Overall, neurons which have a high information content have well separated response distributions to different stimuli.. (**F**) Bar chart showing the number of neurons that were segmented and classified as visually responsive. (**G**) Bar chart showing the number of neurons that had significant information to the local or whole-field motion stimulus sets. (**H**) Average amount of information each neuron contained across fish for the stimulus sets. (**I** & **J**) Histogram of the mutual information across all cells from all fish for the stimulus sets.

### 3.2 An asymmetric distribution of local and whole-field motion encoding neurons along the anterior-posterior axis

One way that there could be regional specialisation within the tectum is if there is a non-uniform distribution in the number of neurons that encode information to the visual stimuli. Therefore, we first wanted to look at the spatial distribution of neurons encoding information to local and whole-field motion. In order to address this we inferred the probability of finding a neuron which encodes information to either local motion or whole-field motion across the anterior-posterior axis of the tectum (Figure 2). To do this we created 10 evenly spaced bins across the anterior-posterior axis of the tectum (Figure 2A). To control for the fact that each bin did not contain an equal number of neurons, we normalised the number of neurons in each bin to the total number of visually responsive neurons in that bin (Figure 2B-D & 2F -H). Neurons which encode information to whole-field motion stimuli were more likely to be found in the anterior regions of the tectum, whilst those which encode information to local motion stimuli were more likely to be found in the posterior regions of the tectum (Figure 2I & J). For each fish, we then calculated an anterior-posterior bias index, which ranges from +1 if all the probability mass is found in the anterior half of the tectum, and -1 if all the probability mass is found in the posterior half of the tectum. Using this index we can see that all fish had an anterior bias for whole-field motion encoding neurons, and a posterior bias for local motion encoding neurons (Figure 2E). Overall, this indicates that there are regional specialisations within the tectum in terms of the spatial distribution of neurons that encode information to visual stimuli.

**Figure 2:**
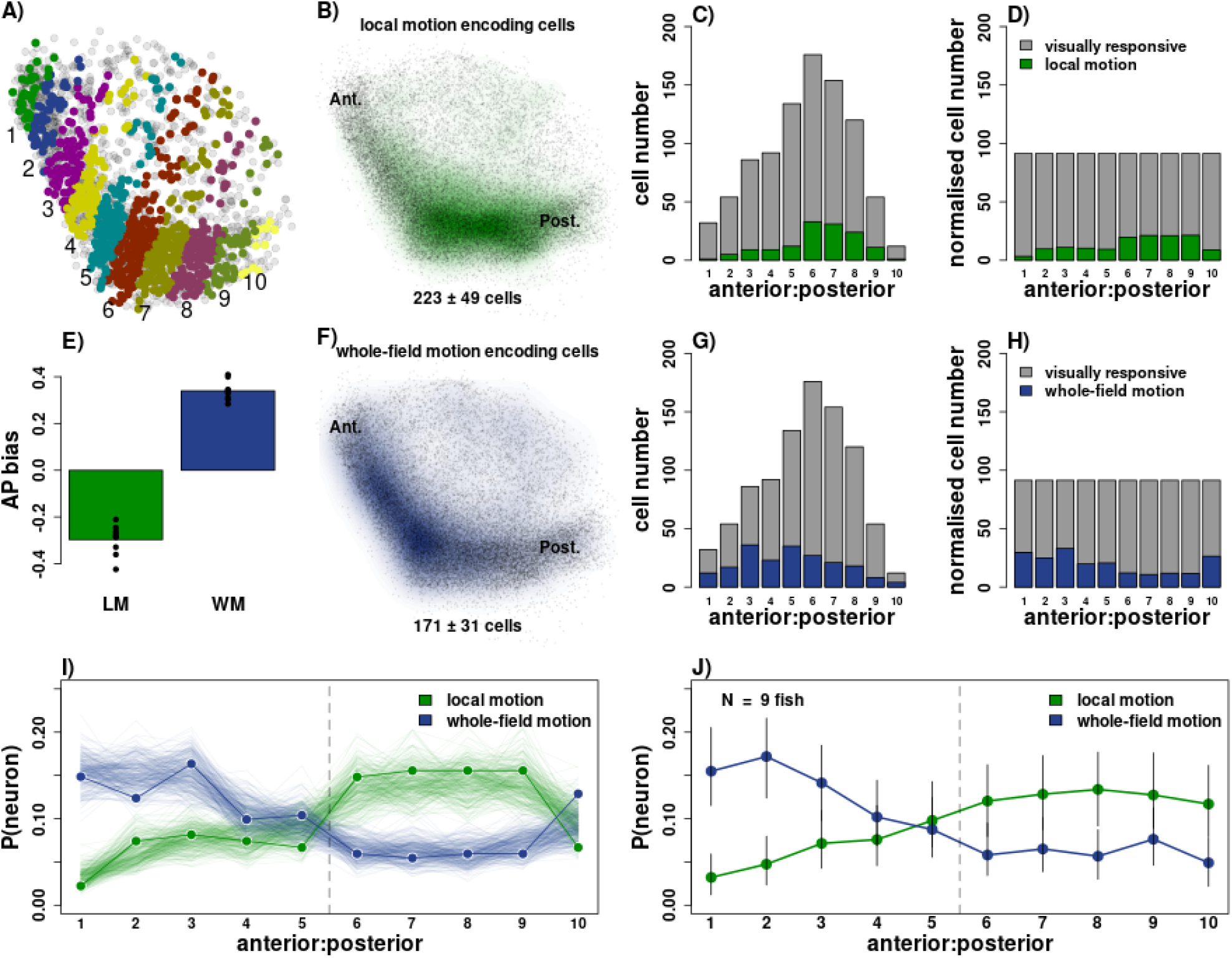
An asymmetric distribution of local and whole-field motion encoding neurons along the anterior-posterior axis of the tectum. (**A**) 10 bins across the AP axis in one example fish, visually responsive neurons colour coded according to bin. (**B**) Probability density map of tectal neurons which encode information to local motion, with the mean number of cells (± sd) per fish underneath, N = 9 fish. (**C**) Total number of visually active neurons (grey) and number of neurons encoding information to local motion (green) per bin across the AP axis in one example fish. (**D**) Number of visually active neurons (grey) and neurons encoding information to local motion (green) after normalisation (methods). (**E**) Mean anterior-posterior bias index for local and whole-field motion encoding cells across fish, each point shows the anterior:posterior bias for a given fish. Positive numbers indicate an anterior bias. (**F**) Probability density map of tectal neurons which encode information to whole-field motion, with the mean number of cells (± sd) per fish underneath. (**G**) Number of visually active neurons (grey) and neurons encoding information to whole-field motion (blue) per bin across the AP axis in one fish example fish. (**H**) Number of visually active neurons (grey) and neurons encoding information to whole-field motion (blue) after normalisation (methods). (**I**) The probability of finding a neuron which encodes information to local or whole-field motion along the anterior-posterior axis of one example fish. The circles show the empirical probability and the lines show draws from the posterior probability of the hierarchical multinomial model (methods). (**J**) The same as in (**I**) but the points show the maximum a posteriori (MAP) estimate of the inferred population (prior) distribution estimated using 9 fish, with error bars showing 95% credible intervals.

### 3.3 The tectum is split into subdomains which preferentially encode local and whole-field motion

A second way that there could be regional specialisation within the tectum is if the amount of mutual information each neuron encodes changes systematically depending on its location within the tectum. For example, do the local motion encoding neurons in the posterior region of the tectum encode more, less, or the same number of bits of information, per neuron, as the neurons in the anterior region of the tectum?

To investigate how the amount of information changes across the tectum we performed 2D Gaussian process regression (Figure 3). We found that for local motion the distribution of information was not uniform within the tectum. In particular, there was a peak in the average amount of information each neuron carries in the tectum’s more posterior and medial regions. One example fish is shown in Figure 3A -H alongside averages of the regression over all fish (Figure 3I & J). Conversely, for neurons which encode information to whole-field motion stimuli there appears to be a more uniform distribution of information across the tectal axes (Figure 3E & H). However, when averaged over fish there does seem to be a peak in the average amount of information in the more anterior regions of the tectum (Figure 3J). Visualising the outputs of the regression in 2D space, along both the anterior-posterior and medial-lateral axes simultaneously, highlights that the posterior-medial region of the tectum encodes high amounts of information to local motion stimuli (Figure 3L), whilst the distribution of information to global motion seems to be more uniform, with a slight bias to the more anterior regions (Figure 3M).

**Figure 3:**
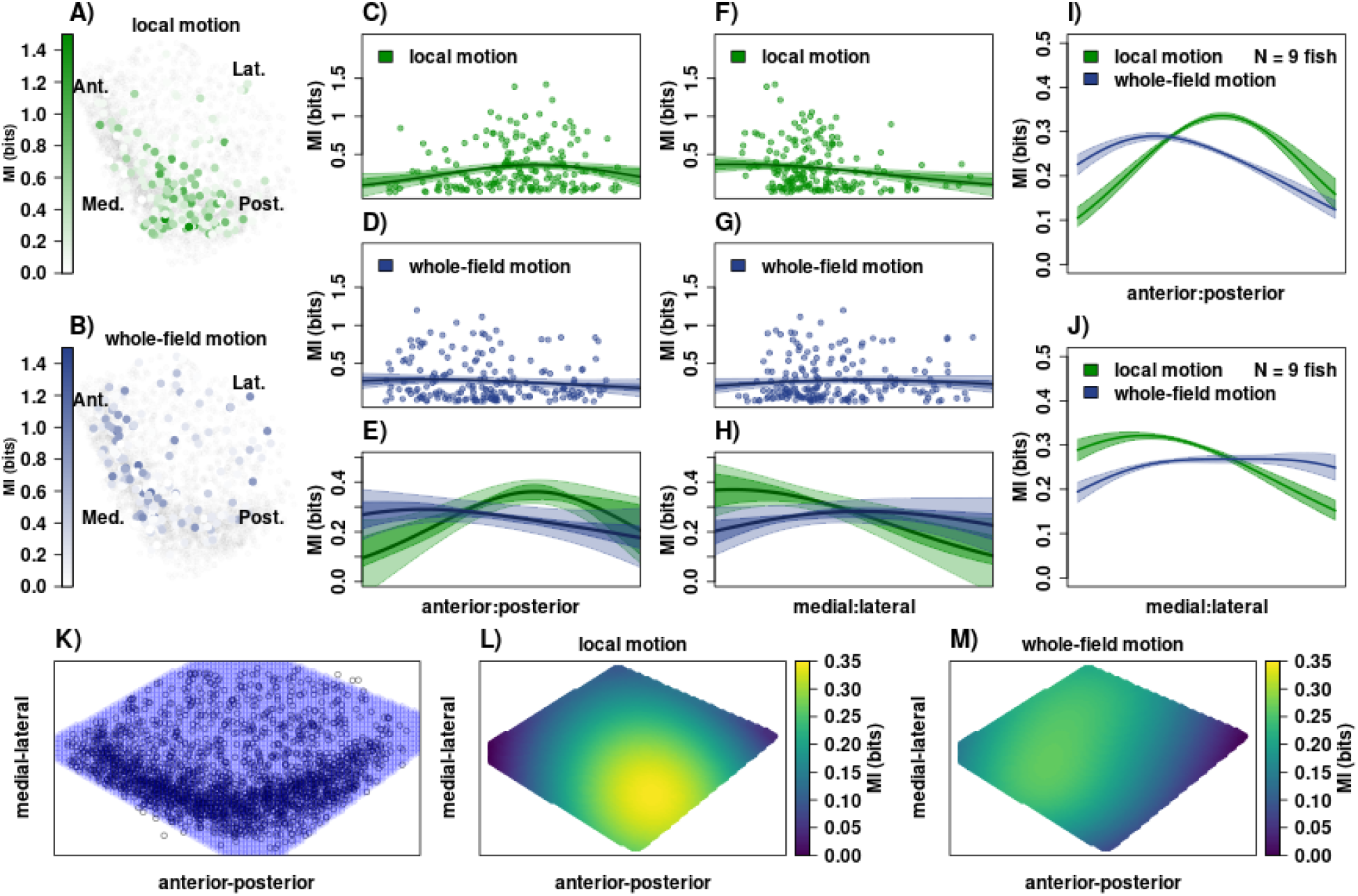
Subdomains within the tectum preferentially encode information to local and whole-field motion. (**A** & **B**) The location of neurons within the tectum that encode information to (**A**) local motion and (**B**) whole-field motion, colour coded according to the amount of information each neuron encodes. (**C** -**E**) Marginal distributions of the Gaussian process regression along the anterior:posterior axis of one example tectum for (**C**) local and (**D**) whole-field motion stimuli. (**E**) shows a zoomed in version of the estimated mean MI. Darker shading represents the 95% credible interval of the mean. Lighter shading represents the estimated standard deviation. (**F**-**H**) Same as in (**C** -**E**) but showing the marginal distribution of the Gaussian process regression along the medial:lateral axis. (**I**) Marginal distribution of the Gaussian process regression along the anterior:posterior axis averaged across 9 fish. (**J**) Marginal distribution of the Gaussian process regression along the medial:lateral axis averaged across 9 fish. (**K**) Region of interest over which to perform the Gaussian process regression. (**L** & **M**) Predicted mean mutual information for (**L**) local and (**M**) whole-field motion across the tectum averaged across 9 fish.

In summary, we see that there can be regional specialisation within the tectum in two ways, 1) a non-uniform distribution in the number of tectal neurons that encode information to the visual stimuli and 2) regional changes in terms of how much information each neuron encodes about the visual stimuli.

### 3.4 Determining functional subtypes

Whilst information theory provides a useful way to quantify how well a neuron can discriminate between groups of stimuli. It doesn’t tell us which features of the stimuli a neuron can discriminate between. For example, a neuron which encodes information about local motion may be able to discriminate between the direction of the local motion stimuli but not the size, or vice versa. Therefore, in order to understand which features of the visual stimuli the neurons are encoding we want to classify the neurons based on the features they could discriminate between. In general, a neuron can be thought of as being able to discriminate a stimulus if its response distribution to that stimulus doesn’t overlap with its response distributions to the other stimuli (Figure S2). To classify the neurons in this manner we first found all the possible combinations of the stimuli which a neuron could in theory discriminate between, given our set of stimuli (Figure S4 and Table S1). Many of these combinations correspond to commonly defined neuronal subtypes such as direction or size selective subtypes. We will therefore refer to these combinations as ‘subtypes’. We next calculated the likelihood of a neuron belonging to each of these subtypes and assigned the neuron to the subtype for which it has the highest likelihood (methods). This gave 22 functional subtypes in total (11 local motion encoding subtypes and 11 whole-field motion encoding subtypes; see Figure S4 for a diagrammatic representation of the subtypes).

Figure 4 shows an overview of the subtypes identified. The subtypes can be split up into four main groups based on the types of features the neurons discriminate between. The first group (Figure 4A; group 1: subtypes 1-4) can best discriminate one specific stimulus from the other three stimuli. The second group (Figure 4A; group 2: subtypes 5-6) can best discriminate between one feature of the stimuli, either direction of motion (subtype 5) or size (subtype 6). These subtypes have been further subdivided based on their direction or size preference. The third group (Figure 4A; group 3: subtypes 7-10) are best able to discriminate between three subsets of the stimuli. For example, the neurons in subtype 10 can discriminate between the direction of motion of the stimuli, but can also discriminate between the size of the stimuli moving in the caudal-to-rostral direction of motion. The fourth group (Figure 4A; group 4: subtype 11) cannot easily discriminate between a particular stimulus, meaning their response distributions to the different stimuli are overlapping. This is also the group with the most number of neurons, which is in keeping with previous findings that the zebrafish optic tectum only contains a small number of well defined feature detectors (Bianco and Engert 2015) and the fact that many visually responsive neurons have a low signal-to-noise ratio (Stringer et al. 2021). There is also one final subtype that we looked for in the analysis, those neurons whose responses can discriminate between all four stimuli (subtype 12 in Figure S4). However, across all 9 fish only two neurons, one for local motion, and the other for whole-field motion, were found (data not shown).

**Figure 4:**
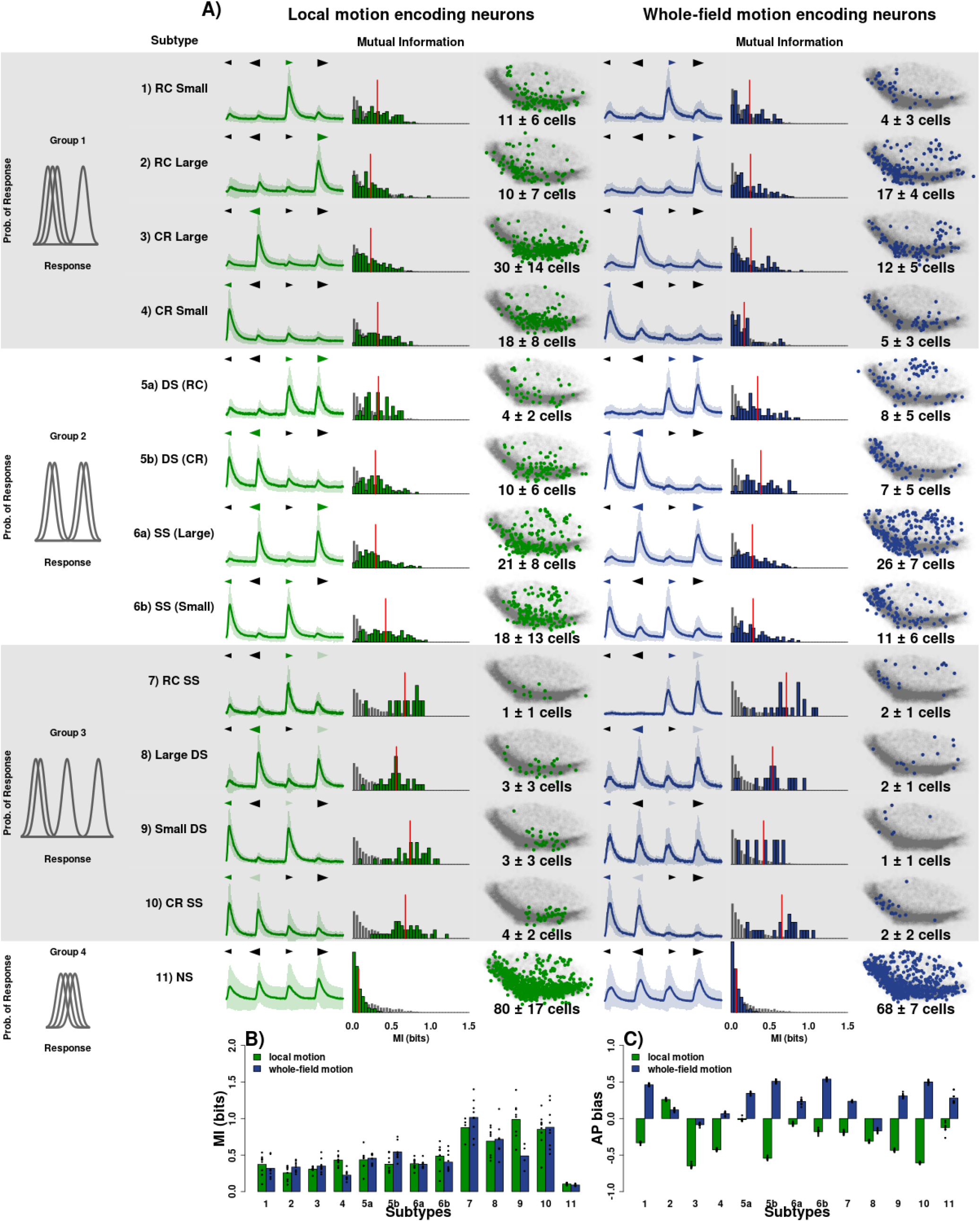
Functional neuronal subtypes. (**A**) Overview of the neuronal subtypes. The left-hand column shows how the subtypes can be split into 3 overarching groups. Group 1 can distinguish one stimulus from the other three. Group 2 can distinguish between two subsets of the stimuli, either direction or size. Group 3 can distinguish between three subsets of the stimuli. Each row shows the average calcium transients, histogram of the mutual information, and spatial localisation of neurons within a subtype. The red bar indicates the mean mutual information within that subtype. The grey histogram shows the information across all neuronal subtypes. The numbers represent the mean number of neurons per subtype across fish (± sd). (**B**) The mean anterior:posterior bias index for the neuronal subtypes. Each point shows the anterior:posterior bias index for a given fish. (**C**) The mean amount of information for each neuronal subtype across fish. Each point shows the average information within a subtype for a given fish. RC: rostral-caudal, CR: caudal-rostral, DS: direction-selective, SS: size-selective, NS: non-selective

The averaged calcium transients (Figure 4A) give an indication that the neurons have been classified correctly. For example, the average calcium transients for subtype 5a show a clear preference for the rostral-to-caudal direction of motion. However, this is an imperfect representation of the subtypes since the transients represent averages over repetitions and neurons, meaning the variance of the responses to the stimuli is necessarily lost. It is not possible to give a perfect population-level overview of the subtypes, however, single-neuron examples can be seen in Figure S5 and Figure S6 for local and whole-motion subtypes, respectively.

The advantage of this approach to classifying neurons is exemplified by the presence of neurons in group 3. It is only possible to detect these subtypes when taking into account the variability of a neuron’s response to each stimulus; something which cannot be done when simply averaging neuronal responses over trials. Whilst there are only a small number of these neurons, they tend to be highly spatially localised to specific regions of the tectum (Figure 4A). Furthermore, defining neuronal subtypes in this manner directly relates to the concept of mutual information. This means we can characterise the subtypes not only based on which features they encode, but also on how much information is encoded. For example, group 3 neurons encode the most amount of information on average, due to their ability to discriminate the most number of visual features, whilst group 4 neurons encode the least amount of information since they are unable to discriminate very well between any of the stimuli (Figure 4B).

Overall we have managed to classify neurons into functional subtypes based on which features of the stimuli they are able to discriminate between. This means we can now relate how well a neuron can discriminate between the stimuli (the mutual information) with the specific features of the stimuli that the neuron is encoding (the subtype).

### 3.5 A division along the anterior-posterior axis is a general organising principle for neurons that encode information about local and whole-field motion

After classifying the neurons into subtypes we next asked whether the asymmetric distribution of local and whole-field motion encoding neurons along the anterior-posterior axis (Figure 2) is due to a particular subtype or subtypes? To do this we calculated the anterior-posterior bias index for each subtype (Figure 4C). As can be seen most local motion subtypes show a posterior bias, whilst most whole-field motion subtypes show an anterior bias, indicating there is a general organising principle for neurons encoding information about local and whole-field motion to be segregated along the anterior-posterior axis (Figure 4C). However, the extent of this bias does vary between subtypes, with some subtypes showing a strong bias e.g subtype 5b and 10, and others showing less bias e.g subtype 11.

We can see the segregation along the anterior-posterior axis particularly well when comparing the distributions of some corresponding local and whole-field motion subtypes. For example, in subtypes 1, 5b, 6b, and 10 there is little spatial overlap between the local motion neurons and corresponding whole-field motion neurons, particularly amongst the periventricular neurons (PVNs) (Figure 5). This indicates that neurons which have the same direction or size selectivity can be localised to different regions of the tectum depending on whether they encode information about local or whole-field motion.

**Figure 5:**
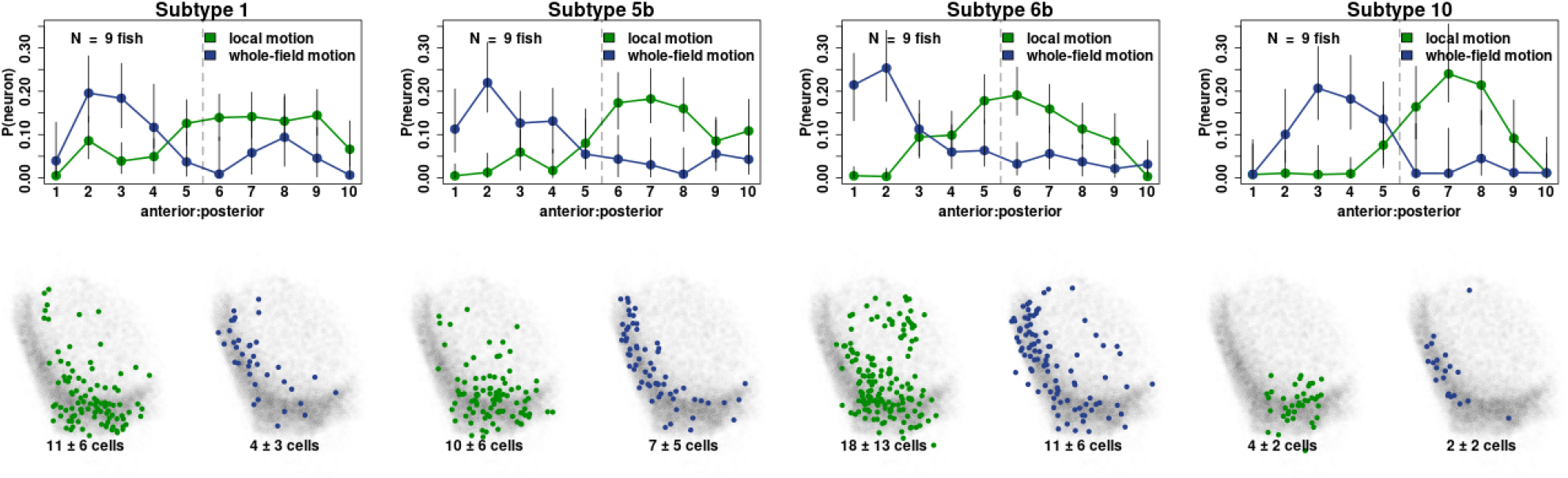
Local and whole-field motion subtypes form opposing spatially localised subdomains. The top row shows the estimated probability of finding a neuron along the anterior-posterior axis of the tectum. Error bars show 95% highest density credible intervals. Bottom row shows the spatial localisation of neurons within the tectum. Numbers show the mean number of neurons within a subtype across fish (± sd).

Previous research has also demonstrated functional segregation along the anterior-posterior axis of the tectum (Förster et al. 2020). In particular, they found a size-dependent mapping for neurons which respond to local motion stimuli, with small size responsive neurons localised to the more posterior regions, and large size responsive neurons localised to the more anterior regions (Förster et al. 2020, their Figure 6). We see a similar pattern in our comparable subtypes; local motion subtype 6b (small) has a greater posterior bias compared to local motion subtype 6a (large) (Figure 4C). However, we also see that large size responsive neurons can be localised to a more posterior region if they respond specifically to caudal-rostral direction of motion (subtype 3; Figure 4C), indicating that the size dependent mapping along the anterior-posterior axis is not preserved for direction-selective neurons.

**Figure 6:**
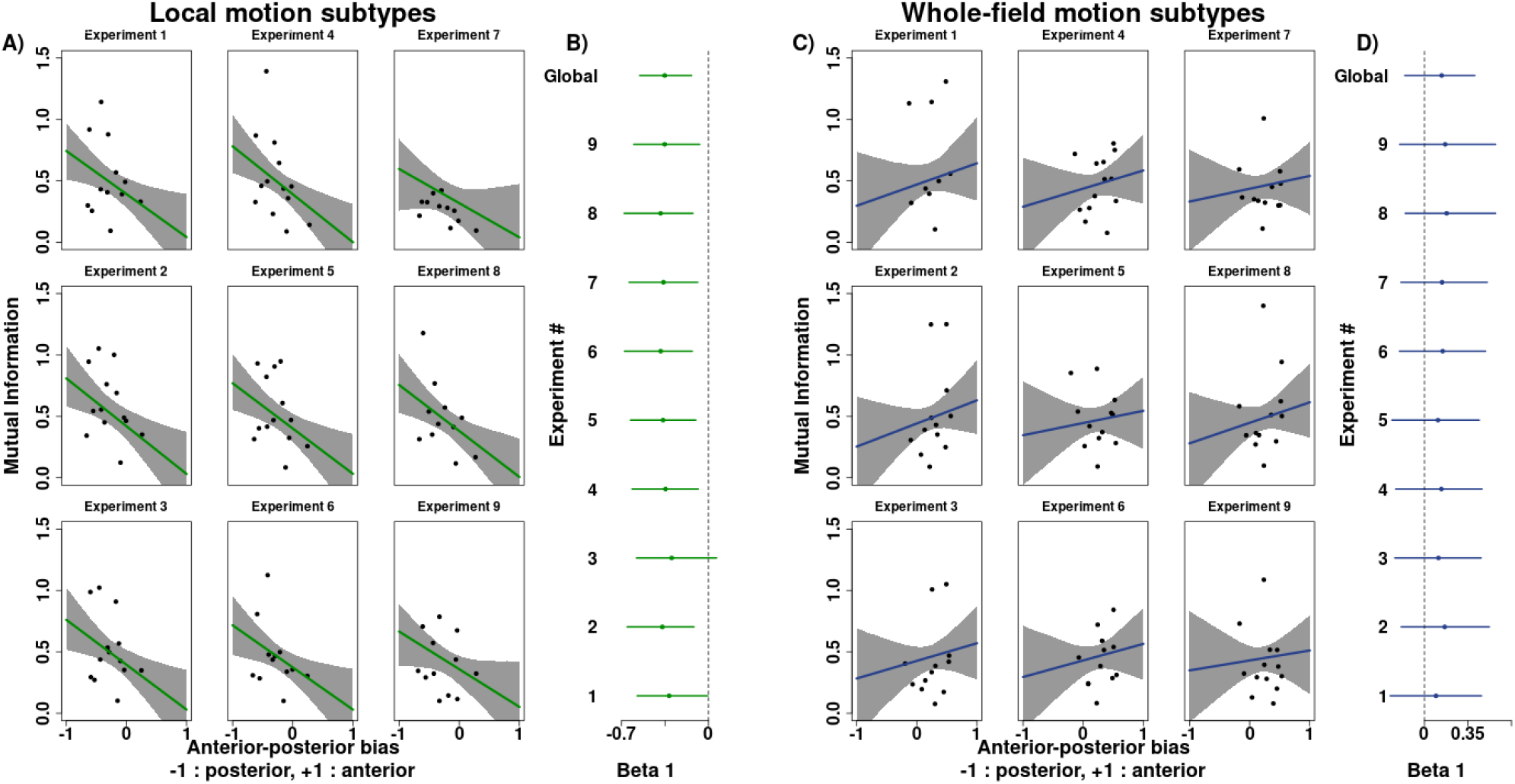
The spatial localisation of subtypes can explain systematic changes in information across the tectum. (**A**) Linear regression between a local motion subtype’s anterior-posterior bias and its mean mutual information. The panels show the 9 fish. Each dot represents a subtype and the shading shows the 95% credible interval of the regression line. (**B**) Estimated *β*_1_ (slope) coefficient for each fish and the global slope coefficient. Error bars show 95% highest density interval. Dotted line shows *β*_1_ = 0. (**C** & **D**) Same as in **A** and **B** but showing whole-field motion subtypes.

### 3.6 Systematic changes in the amount of information across the tectum can be explained by spatially localised functional subtypes

Since some subtypes have a larger anterior-posterior bias than others (Figure 4B) and some subtypes encode more information than others (Figure 4C), we next wanted to ask whether the systematic changes in the amount of information across the axes of the tectum (Figure 3) could be explained by the spatial localisation of the subtypes. To address this question we performed hierarchical linear regression between the anterior-posterior bias of each subtype and the average amount of information within that subtype. We find that for local motion subtypes there is a negative correlation between how much of an anterior-posterior bias a subtype has and the amount of information it encodes (Figure 6A & B; local motion: global slope parameter *β*_1_ = −0.35, 95% HDI[−0.55, −0.14], *P* (*β*_1_ *<* 0) = 99.8%), indicating that local motion subtypes which are localised to the posterior region of the tectum encode, on average, more information than subtypes localised to the anterior regions of the tectum. In particular, we can see this in subtypes 10 and 11 which, whilst they contain only small numbers of neurons, are both highly spatially localised to the posterior region of the tectum and each contain neurons which encode large amounts of information. For whole-field motion subtypes, whilst the central estimate for each slope parameter is positive (indicating subtypes localised to the anterior tectum encode higher amounts of information), in every case the 95% highest-denisty interval (HDI) of the parameters extend into negative values indicating a lack of certainty in the slope parameters being positive (Figure 6C & D; whole-field motion: global slope parameter *β*_1_ = 0.14, 95% HDI[−0.16, 0.41], *P* (*β*_1_ *>* 0) = 83.5%), which is in keeping with the idea that the amount of information for global motion is more uniform across the tectal axes (Figure 3).

## 4 Discussion

### 4.1 An emerging view of regional specialisation within the optic tectum

This study aimed to investigate whether there were regional specialisations in how visual information is encoded within the optic tectum of the larval zebrafish, not only in terms of the spatial distribution of neuronal subtypes but also in terms of how well these subtypes could discriminate between the visual stimuli. Our findings are consistent with an emerging view that the tectum contains regions of functional specialisation, rather than a uniform distribution of neuronal subtypes. For example, our finding that there is an over-representation of local motion encoding neurons in the posterior region of the tectum (Figures 2 and 4) is in keeping with the discovery of an over-representation of direction-selective small-dot responsive neurons in the tectum’s more posterior regions (Förster et al. 2020). However, it is not simply the localisation of these neurons to the posterior tectum which causes a regional specialisation. The amount of information those neurons encode also changes systematically, with the posterior neurons encoding higher amounts of information, on average, than those in the anterior (Figure 3 and 6).

Due to the tectum’s topographic mapping, the existence of a posterior specialisation for detecting local motion stimuli implies that there would be changes in how well local motion stimuli are encoded at different points along the visual field. This is supported by the fact that there is an increased ability to decode stimuli in the more temporal regions compared to stimuli in more nasal regions early on in development (4-5 dpf) seemingly due to an increase in the amount of information encoded by neurons localised to the posterior region of the tectum (Avitan et al. 2020). Interestingly, this bias for decoding stimuli in the temporal visual field disappeared over development. However, this was due to neurons in the posterior region of the tectum shifting their receptive fields to more anterior positions, rather than the development of local motion encoding neurons in the anterior tectum. Furthermore, a study which analysed tectal activity in 6-8 dpf freely moving fish during prey capture observed that increasing numbers of neurons were activated as prey stimuli were localised to more temporal regions of the visual field (Marques et al. 2020). Due to the topographic mapping between prey location and neuronal activation, this also resulted in a denser activation of neurons in the more posterior regions of the tectum, in keeping with our own findings. Another study which analysed tectal activity in freely moving larvae also found that there was a posterior bias in tectal cell activity when the larvae were presented with live prey (Muto et al. 2013). However, the authors also found that tectal activity immediately preceding prey capture behaviour (defined by eye convergence) was shifted into a more anterior position. This indicates that posterior tectal activity may play a role earlier on in the initial detection phase of hunting.

For whole-field motion encoding neurons, we find that whilst the amount of information each neuron encodes is relatively uniform across the tectum, there is an anterior bias in terms of the number of neurons encoding whole-field motion. As far as we are aware, this has not been reported before. Whilst it may seem counter-intuitive for neurons that respond to a global signal to be localised to a particular region of the tectum, previous studies have demonstrated many animals show a bias in terms of the regions of visual space they use to estimate optic flow (Alexander et al. 2022, Fujimoto and Ashida 2019, Krapp and Hengstenberg 1996, Nityananda et al. 2017). This is thought to be due to variations in the statistics of the visual scene, which means some regions of visual space provide more accurate information about optic flow than other regions. Zebrafish larvae are thought to interpret whole-field gratings as experiencing optic flow due to the ability of wholefield gratings to initiate stabilisation reflexes such as the optomotor response and optokinetic response (Brockerhoff et al. 1995, Kubo et al. 2014, Naumann et al. 2016). Therefore, it might not be so surprising that we see regional changes in the density of whole-field motion encoding neurons within the tectum if particular portions of the visual scene provide better contextual information about optic flow.

### 4.2 Information encoding in the tectum

We find that the majority of visually responsive neurons only provide a small amount of information regarding the visual stimuli (Figure 1I & J) and don’t fit neatly into a particular feature detector subtype (Figure 4; subtype 11). This is in line with previous research in zebrafish (Bianco and Engert 2015, Chen et al. 2018), and is in keeping with the idea that population activity over groups of neurons provides a way to mitigate single-neuron noise (Faisal et al. 2008). This, however, highlights a limitation of the current study; we have only calculated the mutual information on single neurons, without taking into account the joint activity across neighbouring neurons. In the future it would be interesting to see how taking into account the population activity affects the regional specialisations we have seen within the tectum. Indeed, it is known that the structure of the population activity, such as noise correlations between neurons, can affect the ability of the population to encode information (Azeredo Da Silveira and Rieke 2021, Kohn et al. 2016, Panzeri et al. 2022). Although, the extent to which these noise correlations affect the encoding capacity of the population is not well established (Stringer et al. 2021). Unfortunately, this is something that cannot be assessed in the current study due to the inherent difficulties in calculating the information content over large populations of neurons. However, recently developed methods have been used to look at population encoding in the mouse cortex, and provide a framework for how to approach such a problem (Kafashan et al. 2021, Rumyantsev et al. 2020, Stringer et al. 2021). Indeed, in the future, the zebrafish tectum may provide a particularly useful model to study how the structure of the population activity affects the encoding of information, due to its relatively small size and optical accessibility.

## 5 Materials and Methods

### 5.1 Zebrafish maintenance

Zebrafish were maintained at 28.5°C on a 14 h ON/10 h OFF light cycle in Danieau solution (58mM NaCl, 0.7 mM KCl, 0.4 mM MgSO4, 0.6 mM Ca(NO3)2, 5 mM HEPES, pH 7.6). Transgenic line used: Tg(HuC:H2B-GCaMP6s; casper). Sex of individual animals not known. Larvae were fed live rotifers daily from 5 dpf and were raised in petri dishes over a bed of gravel. This work was approved by the local Animal Care and Use Committee (King’s College London), and was carried out in accordance with the Animals (Experimental Procedures) Act, 1986, under license from the United Kingdom Home Office.

### 5.2 Imaging

Non-anesthetized 7 dpf Tg(HuC:H2B-GCaMP6F; casper) larvae were immobilized at in 2% low melting point agarose (Sigma-Aldrich) prepared in Danieau solution and mounted dorsal side up in a custom-made perspex cylindrical Danieau-filled chamber. Fish were positioned such that the right eye was 20 mm away pointing towards a semi-circular screen covered in a grey diffusive filter which occupied 153° x 97° of visual azimuth and elevation. Visual stimuli were projected using a P2JR pico-projector (AAXATech). To avoid interference of the projected image with the signal collected by the detector, a red long-pass filter (Zeiss LP590 filter) was placed in front of the projector. To prevent drift of the larvae through the agarose during the experiment, the larvae were allowed to settle for 2 hours prior to recording.

Functional imaging was conducted using a custom built 2-photon microscope (Independent NeuroScience Services). Excitation was provided by a Mai Tai HP ultrafast Ti:Sapphire laser (Spectraphysics) tuned to 940nm. Emitted light was collected by a 16x, 1 NA water immersion objective (Nikon) and detected using a gallium arsenide phosphide (GaAsP) detector (ThorLabs). Images (256 x 256 pixels) were acquired at a frame rate of 50 Hz by scanning the laser in the x-axis with a resonant scanner and in the y-axis by a galvo-mirror. The focal plane was adjusted in 10 *µ*m steps using a piezo lens holder (Physik Instrumente). This allowed for volumetric data consisting of 7 focal planes to be collected at a volume rate of 7.28 Hz. Scanning and image acquisition were controlled by Scanimage Software (Vidrio Technologies).

### 5.3 Visual stimuli

Visual stimuli were custom written in C++ using the opencv package. Visual stimuli used in the present analysis include dots covering 5° and 26° of visual angle moving forwards (caudal to rostral) and backwards (rostral to caudal) along the visual azimuth at a speed of 25°/s. These were dark dots presented over a mean grey background. Sinusoidal gratings with a spatial frequency of 1/5° and 1/26° (which correspond to wavelengths covering 5° and 26° of visual angle) and temporal frequency of 25°/s also moving forwards (caudal to rostral) and backwards (rostral to caudal). The presentation of the dots and gratings stimuli were conducted using 6 second long epochs, followed by a 25 second blank screen to allow the calcium transients to return to baseline. Stimuli faded in at the beginning of each epoch and each visual stimulus was presented a total of 10 times in a pseudo-randomised order.

### 5.4 Preprocessing functional imaging data

Visually evoked functional imaging data was aligned and segmented using the Suite2p Python package (Pachitariu et al. 2017). Only segments within the tectum with a probability *>* 0.5 of being a cell were used for further analysis. For each neuronal calcium trace, a baseline which corrects for low-frequency drifts was calculated using a previously described method (Diana et al. 2019). Briefly, a hidden-Markov model decomposes the neural calcium activity into the sum of calcium transient (*c*_*k*_), baseline activity (*b*_*k*_) and a source of Gaussian noise. The hidden on/off state of a neuron is modelled as a Bernoulli process, giving rise to time points of inferred neural activity. The calculated background activity was subtracted from each mean normalised neuron.

To classify a neuron as being visually responsive or not, the binarised activity vectors from the hidden markov model were used to calculate a correlation coefficient between time points when a neuron was active and timepoints when a stimulus was being presented: Let *x*_*i*_ be the binarised neuronal activity at time point *i* (1 = active, -1 = not active) and *s*_*i*_ be the visual stimulus vector at timepoint *i* (1 = visual stimulus being presented, -1 = no visual stimulus being presented). The correlation coefficient is given by:

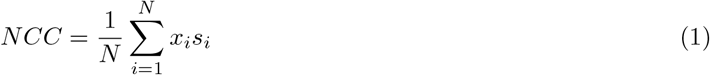

where N is the number of frames. A neuron was classified as visually responsive if its correlation was 0.02 standard deviations higher than expected given the neuron was firing randomly:

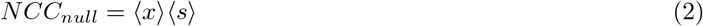

where ⟨⟩ represents the average value of the binarised activity vector or stimulus vector.

### 5.5 Aligning fish to a standard space

The spatial coordinates of neurons across different fish were aligned to a common reference frame to allow a direct comparison of the spatial distribution of neurons across different experiments. This was performed using the Advanced Normalisation Tools (ANTs) software (Tustison et al. 2014). First, for each experiment, a reference stack of the tectum was taken on the 2-photon using the same xy coordinates as the functional imaging stack, but with a 2*µ*m step in z. This stack covered the whole dorso-ventral region of the tectum with 200 slices per stack. The functional imaging planes were then aligned to their corresponding planes in the reference stack using the SyN method via the ANTsR package. These functional imaging planes were then swapped into the reference stack. These reference stacks were then aligned across experiments, choosing one fish to be the ‘standard’ fish, again using the SyN method. For both alignment steps the transformations were applied to the coordinates of the cell bodies from suite2p.

### 5.6 Calculating the mutual information

The mutual information between the stimulus and response of a neuron is given by:

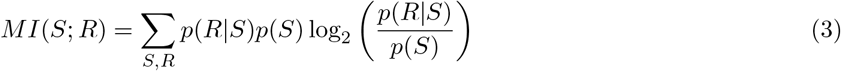

Where *P* (*S*) is the probability of showing a particular stimulus and *P* (*R*|*S*) is the probability of response to each specific stimulus. The probability of a particular stimulus being shown was kept uniform for all experiments, and since each stimulus set is composed of 4 stimuli, *P* (*S*) = 0.25 across all experiments and stimulus sets.

To estimate *P* (*R*|*S*) for each neuron the mean fluorescence over the epoch was taken for each repetition of a given stimulus. This gives us a distribution of responses for each stimulus. We then modelled *P* (*R*|*S*) as a Gaussian distribution over the repetitions and estimated the parameters using method of moments from the ‘fitdistr’ package.

Using these estimated probabilities we can calculate two information entropies. First, we can calculate the conditional entropy:

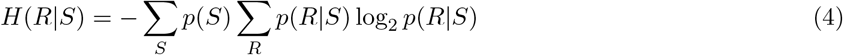

which quantifies the variability of the response to the stimuli. Second, we can calculate the marginal entropy:

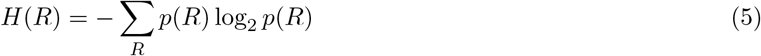

which quantifies the variability of the average response of a neuron. We can use these two information entropies to calculate the mutual information:

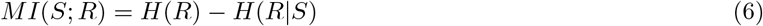

Once we had estimated *P* (*R*|*S*) then averages over the response, as in equations 4 and 5, were calculated using a grid approximation with a grid size of 1000. The interval over which the summation was performed was defined as minus three standard deviations of the smallest estimated mean response, to plus three standard deviations of the largest estimated mean response. For each neuron, the standard deviation was taken to be the largest estimated standard deviation from each of the four stimuli in the set.

Since *P* (*R*|*S*) must be estimated empirically from the data, and limited sampling can lead to a positive bias in the estimation of the mutual information (Treves and Panzeri 1995), it was necessary to apply a correction factor to the estimated mutual information for each neuron. We used a shuffling procedure whereby the stimulus labels across both local and whole-field stimulus sets were randomly allocated to a response, and the mutual information of this shuffled data was calculated. We did this 20 times and calculated the average mutual information across the shuffled data, which was then subtracted from the actual estimated mutual information. Any neuron that still had positive mutual information following this procedure was classified as having significant information to a stimulus set. Only neurons with significant mutual information to a stimulus set were used in the analysis of that set. We also added the constraint that for a neuron to have significant information to one stimulus set (either local or global motion) it must not have significant information to other set.

### 5.7 Multinomial model of neuronal distribution

A hierarchical multinomial model was used to infer the probability of finding a neuron across the anterior:posterior axis of the tectum. First we counted all of the neurons which had significant mutual information to a given stimulus set in each of the bins across the anterior:posterior axis. To control for the fact that the bin sizes were not equal we normalised the cell counts in the following way. First, the number of neurons with significant mutual information in each bin was normalised to the total number of visually active neurons in that bin. This gives a ratio for each bin. Each of these ratios was then normalised to the sum of the ratios across the bins and then multiplied by the total number of neurons with significant information across the bins. This redistributes the neuronal counts based on the size of the bins whilst keeping the total number of neurons the same as the non-normalised count.

We modelled the normalised number of neurons across the 10 bins as coming from a multinomial distribution,

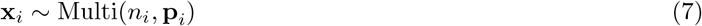

where **x**_*i*_ is a vector representing the 10 bins, with each element containing the normalised number of neurons in a bin for fish *i, n*_*i*_ is the total normalised number of neurons across bins for fish *i*, and **p**_*i*_ is the vector of probabilities of finding a neuron in each bin for fish *i*. We can place a Dirichlet prior over **p**_*i*_,

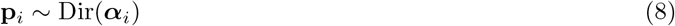

where ***α***_*i*_ is a vector of positive reals, with one entry per bin. Since we are trying to infer **p** across each fish, the data can therefore be considered in a hierarchical manner, where ***α***_*i*_ is assumed to come from an underlying global distribution across all fish. We can therefore place a uniform hyperprior over ***α***_*i*_,

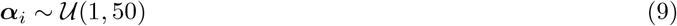

Inference for the model defined in equations 7 to 9 was performed by Markov chain Monte Carlo (MCMC) sampling using the JAGS (Plummer 2003) probabilistic programming language via the rjags interface.

### 5.8 Gaussian process regression model

To infer the amount of information across the axes of the tectum we used a Gaussian process regression model. We can think of the amount of mutual information, *z*, across the xy plane of the tectum, **x**, being defined by some unknown, latent function, *f*, such that:

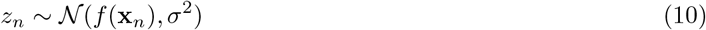

That is, the amount of information at any given point in the tectum is normally distributed with mean, *f* (**x**), and noise *σ*^2^, which is assumed to be i.i.d over each observation, *n*. This defines the likelihood of our data. To minimise our assumptions about *f* (**x**) we can specify a prior over *f* in terms of a Gaussian process. A Gaussian process is parameterized by two functions: a mean function *m*(**x**) and covariance function *k*(**x, x**^*′*^).

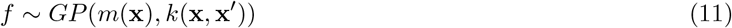

In our model we have set *m*(**x**) = 0. In general, this can be done without loss of performance over the domain for which we have data, since the posterior is able to capture the mean in this range.

We define the covariance function as the exponentiated quadratic function:

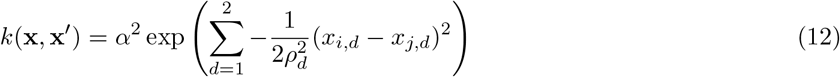

where *d* represents the dimensionality of the input space, which in our case is the 2-dimensional input of the xy spatial coordinates of the neuron. Here, *α* and *ρ*_*d*_ are hyperparameters to be inferred, with *α* specifying the marginal deviation of the function values, i.e the uncertainty around the mean at a given point. The 2 parameters *ρ*_1_ and *ρ*_2_ each specify the length scales across each of the two spatial dimensions in the xy plane. Each *ρ* determines how quickly the covariance between neighbouring points decreases in either dimension. Using the exponentiated quadratic function implies that the underlying mean mutual information changes smoothly across the tectum, with a higher degree of dependence between nearby neurons.

We define the hyperparameters and the noise parameters collectively as ***θ*** = {*α, ρ*_1_, *ρ*_2_, *σ*}

The use of a Gaussian process prior makes the assumption that, for any given set of xy coordinates, **x**, the mean mutual information at those points is jointly Gaussian with mean *m*(**x**) and covariance *k*(**x, x**^*′*^):

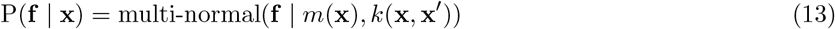

### 5.9 Gaussian process regression inference

Given the above model, and a set of data points 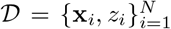, where **x** is the location of the neurons in the tectum, and **z** is the amount of information each neuron encodes, we want to predict the mean mutual information at any point in the tectum, *f* (**x**). We first define a grid across the xy plane of the tectum: **x**_*∗*_ = {**x**_*∗***1**_, …, **x**_*∗***n**_}, at points where we want to evaluate the latent function, *f*, where **f**_*∗*_ = {*f* (**x**_*∗***1**_), …, *f* (**x**_*∗***n**_)} are the corresponding latent function values. We can then define the joint distribution of the measured mutual information values from the data **z**, and the value of the latent function at the predicted points **f**_*∗*_, as:

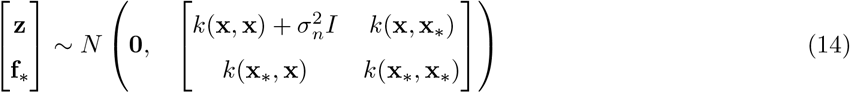

Since **z** and **f**_*∗*_ are jointly Gaussian random vectors, we can derive the predictive distribution **f**_*∗*_ | **x**_*∗*_, **z, x** as:

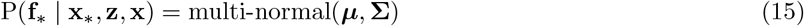

where,

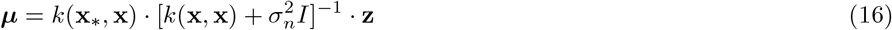

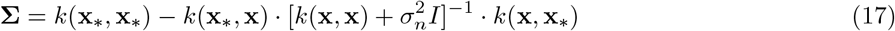

Equation 15 represents the posterior distribution of the latent function values i.e. the probability of the mean mutual information across the tectum. We have, however, so far only considered the case where the values for the hyperparameters, ***θ*** are known. In general we do not know what values these hyperparameters should take. To incorporate our uncertainty about these values, we can integrate over ***θ*** in the following manner:

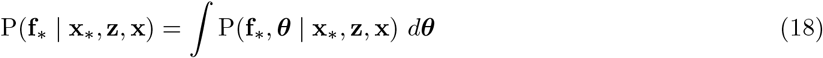

The integral can be decomposed into two terms:

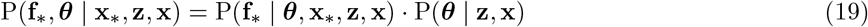

where the first term on the right corresponds to equation 15 for a given set of hyperparameter values, and the second term corresponds to the posterior distribution of the hyperparameters. To evaluate this posterior distribution we can use Markov Chain Monte Carlo (MCMC) methods.

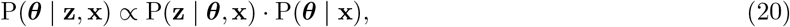

where P(**z** | ***θ*, x**) corresponds to the marginal likelihood of the data:

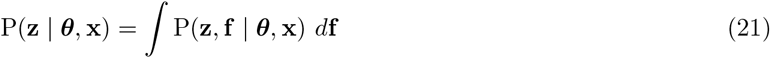

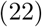

From (14) we can see that this is equal to,

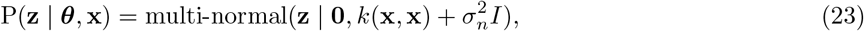

and P(***θ*** | **x**) in (20) corresponds to the prior probability of the hyperparameters. In the analysis, we used the following priors:

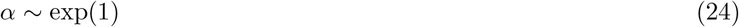

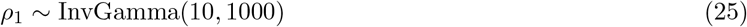

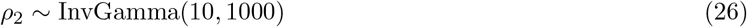

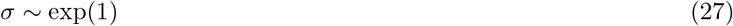

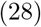

To produce draws from the posterior using MCMC methods we used Stan (Stan Development Team 2012) version 2.26.13 via the RStan (Stan Development Team 2020) interface. For each Gaussian Process, 8000 iterations were split on 4 parallel chains each with a warm-up of 1000 iterations.

### 5.10 Classifying neuronal subtypes

A functional neuronal subtype is defined by its pattern of activity, often in response to some kind of sensory input. For example, if the activity of a neuron was being recorded in response to two visual stimuli moving in opposite directions, the neuron could be classed as direction selective if it robustly modulated its activity depending on the stimulus direction. However, if the neuron responded in a similar manner to both stimuli, it would be classed as being non-selective. In the direction-selective case, we can think of the neuron’s response as coming from two separate response distributions, one for each stimulus. In the non-selective case, the neuronal response can be thought of as coming from a single response distribution. We can, therefore, more generally define a neuronal subtype as the collection of distinct response distributions elicited by an array of visual stimuli.

Therefore, to pre-define our neuronal subtypes we need to determine all the possible collections of response distributions that a set of stimuli can elicit. This corresponds to determining all possible partitions of the set of stimuli into non-empty subsets. For four stimuli we get a total of 15 possible partitions (Table 1). Each partition can be thought of as a functional subtype, many of which correspond to commonly defined neuronal subtypes (Table 1). Each subset within a partition contains one or more stimuli. These subsets can be thought of as defining a single response distribution for a neuron, so if two stimuli fall within the same subset then the response of a neuron to those stimuli is assumed to come from the same distribution i.e the neuron doesn’t distinguish between those stimuli within a subset. For example, a neuron which is classed as size selective, but not direction selective, can be thought of as having the same response distribution to both small stimuli moving in opposite directions and both large stimuli moving in opposite directions. This would correspond to partition 8 of Table 1. See Figure S3 for an example neuron that is classified as size selective.

**Table 1:**
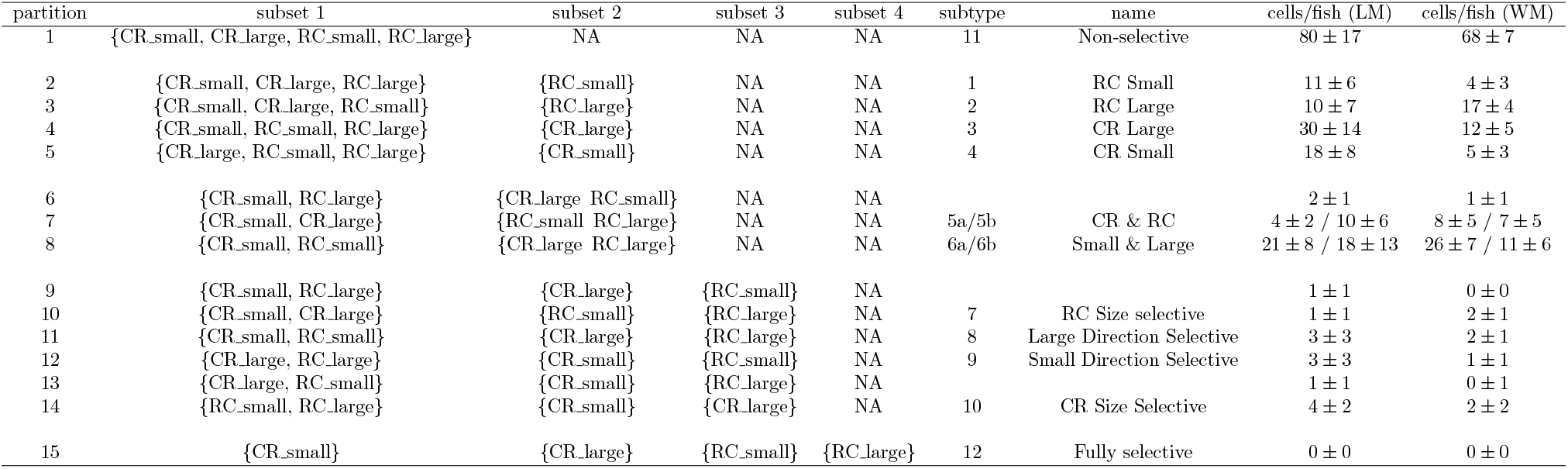
Details of neuronal subtypes. Each partition represents a unique division of the stimuli into non-empty subsets. Partitions 6, 9, and 13 don’t correspond to biologically meaningful subtypes, and due to the small number of neurons that were associated to them, they were removed from further analysis. CR: caudal-rostral, RC: rostral-caudal, LM: local motion, WM: whole-field motion

We, therefore, have 15 possible partitions to which a neuron could be assigned. To assign a neuron to one of these partitions we calculate the marginal log likelihood for each partition and assign the neuron to the partition for which it has the highest log likelihood.

We calculate the marginal log likelihood as follows: for each subset, *n*, within a partition, we take the distribution of responses, *D*_*n*_ (calculated as in section 3.5) for all stimuli within that subset. We assume this distribution of responses within the subset comes from a Gaussian with unknown mean, *µ*, and precision, *τ* :

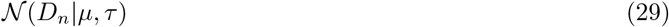

We can define a prior over the parameters as:

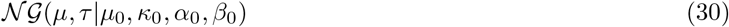

Where 𝒩 𝒢 is a normal-gamma distribution. We can take advantage of the conjugate nature of the prior distribution to analytically derive the marginal likelihood of the data:

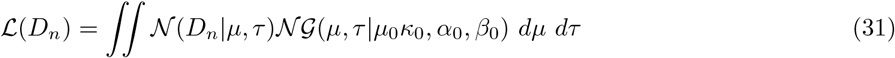

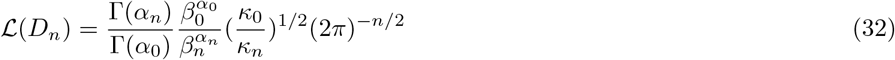

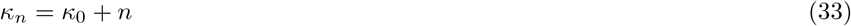

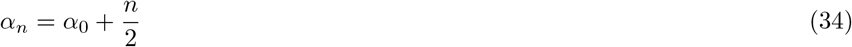

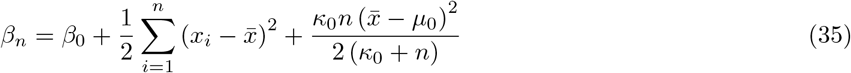

Where *µ*_0_, *κ*_0_, *α*_0_, *β*_0_ are the parameters of the conjugate normal-gamma prior. The priors used in the analysis were:

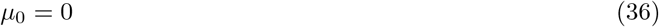

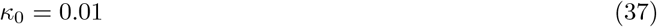

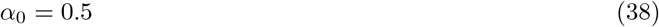

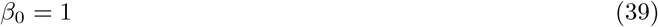

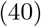

We can, therefore, calculate the log likelihood of a neuron belonging to each partition by summing over each subset, *n*, within a partition:

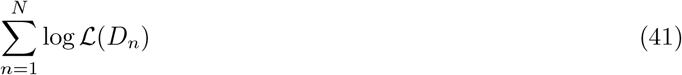

where *N* is the total number of subsets within a partition. We can then associate each neuron to the partition for which it has the highest log likelihood value. See Figure S3 for an example neuron.

Partition 7 defines direction selective neurons. A neuron assigned to this partition responds preferentially to either one of the two directions. The neurons in this partition were further subdivided into two separate subtypes: ‘CR’ for neurons which respond preferentially to the caudal-to-rostral direction, and ‘RC’ for neurons which respond preferentially to the rostral-to-caudal direction. To assign a neuron to either group we calculated a selectivity index as:

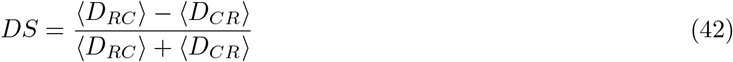

where *D*_*RC*_ is the neuron’s distribution of responses to the RC small and RC large stimuli, and *D*_*CR*_ is the neuron’s distribution of responses to the CR small and RC large stimuli. Plotting the *DS* index across all neurons assigned to partition 7 demonstrates that the neurons within this partition naturally fall into two segregated groups (Figure S7).

The same process was repeated for partition 8 which defines size selective neurons. The neurons in this partition were further subdivided into two separate subtypes: ‘small’ for neurons which respond preferentially to the small size stimuli and ‘large’ for neurons which respond preferentially to the large size stimuli. To assign a neuron to either group we calculated a size selectivity index as:

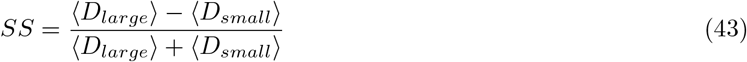

where *D*_*large*_ is the neuron’s response to the RC large and CR large stimuli, and *D*_*small*_ is the neuron’s response to the RC small and CR small epochs. Plotting the *SS* index across all neurons within partition 8 demonstrates that the neurons within this partition naturally fall into two segregated groups (Figure S7).

A similar process was repeated for partitions 2-5. In this case, a selectivity index was calculated as:

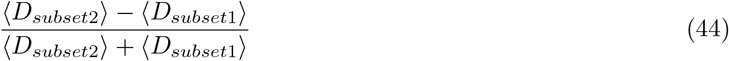

where *D*_*subset*2_ is the neuron’s response to the stimuli in subset 2 and *D*_*subset*1_ is the neuron’s response to the stimuli in subset 1. All positive values indicate neurons that respond preferentially to the stimulus in subset 2. As can be seen in Figure S8 these constitute the vast majority of the neurons in the partition. The small number of neurons with negative selectivity (which responded preferentially to subset 1) were removed from the partition and discarded.

### 5.11 Hierarchical linear regression of subtype location and mutual information

Hierarchical linear regression was used to infer whether there was a relationship between the spatial distribution of a subtype (anterior:posterior bias) and the average amount of mutual information within that subtype. For each subtype, *i*, and each fish, *j*, we assume the average amount of information within a subtype, *M*, is linearly related to it’s anterior:posterior bias along the tectal axis, *B*,

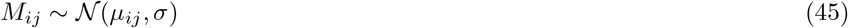

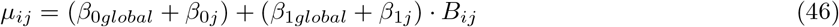

We assume the errors, *σ*, are independent and identically distributed, for which we use an exponential prior.

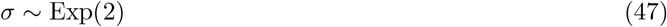

We have also decomposed the intercept, *β*_0_, and slope, *β*_1_, into two components. A global component and a fish-specific component. The priors for the global coefficients are:

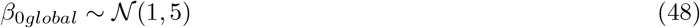

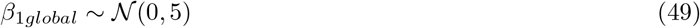

We further assume that the fish-specific coefficients, *β*_0*j*_ and *β*_1*j*_ come from a joint prior distribution,

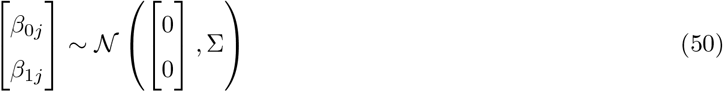

Where Σ is the covariance matrix. To more easily place a prior on the covariance matrix, we can decompose Σ into 3 matrices,

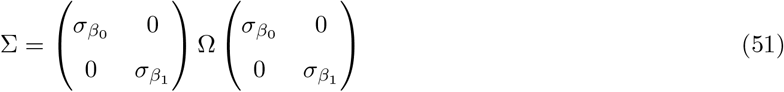

where 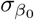 and 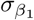 are the standard deviations of *β*_0_ and *β*_1_, respectively and Ω is a correlation matrix. We can then place priors over these parameters,

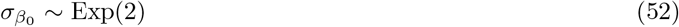

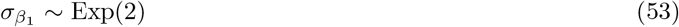

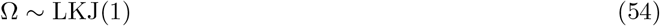

The dependence of *β*_0_ and *β*_1_ on Σ can lead to difficulties in sampling from the posterior. To more efficiently sample from the posterior we can reparameterise the model in the following manner. First, we can sample from two independent standard normal distributions,

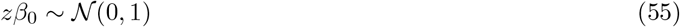

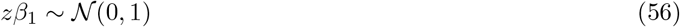

We next want to generate correlated samples of the fish-specific coefficients, according to the inferred co-variance matrix Σ. To do this we can transform our two independent standard normal distributions,

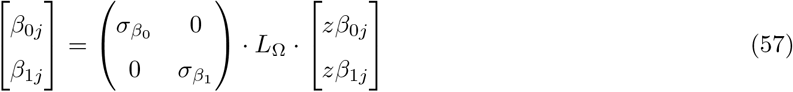

Where *L*_Ω_ is the Cholesky factor of Ω. We can place a prior over *L*_Ω_ such that,

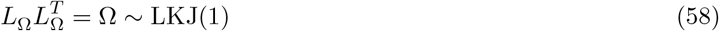

To produce draws from the posterior using MCMC methods we used Stan (Stan Development Team 2012) version 2.26.13 via the RStan (Stan Development Team 2020) interface. We sampled 20000 iterations which were split on 4 parallel chains, each with a warm-up of 2500 iterations.

---

**Figure S1:**
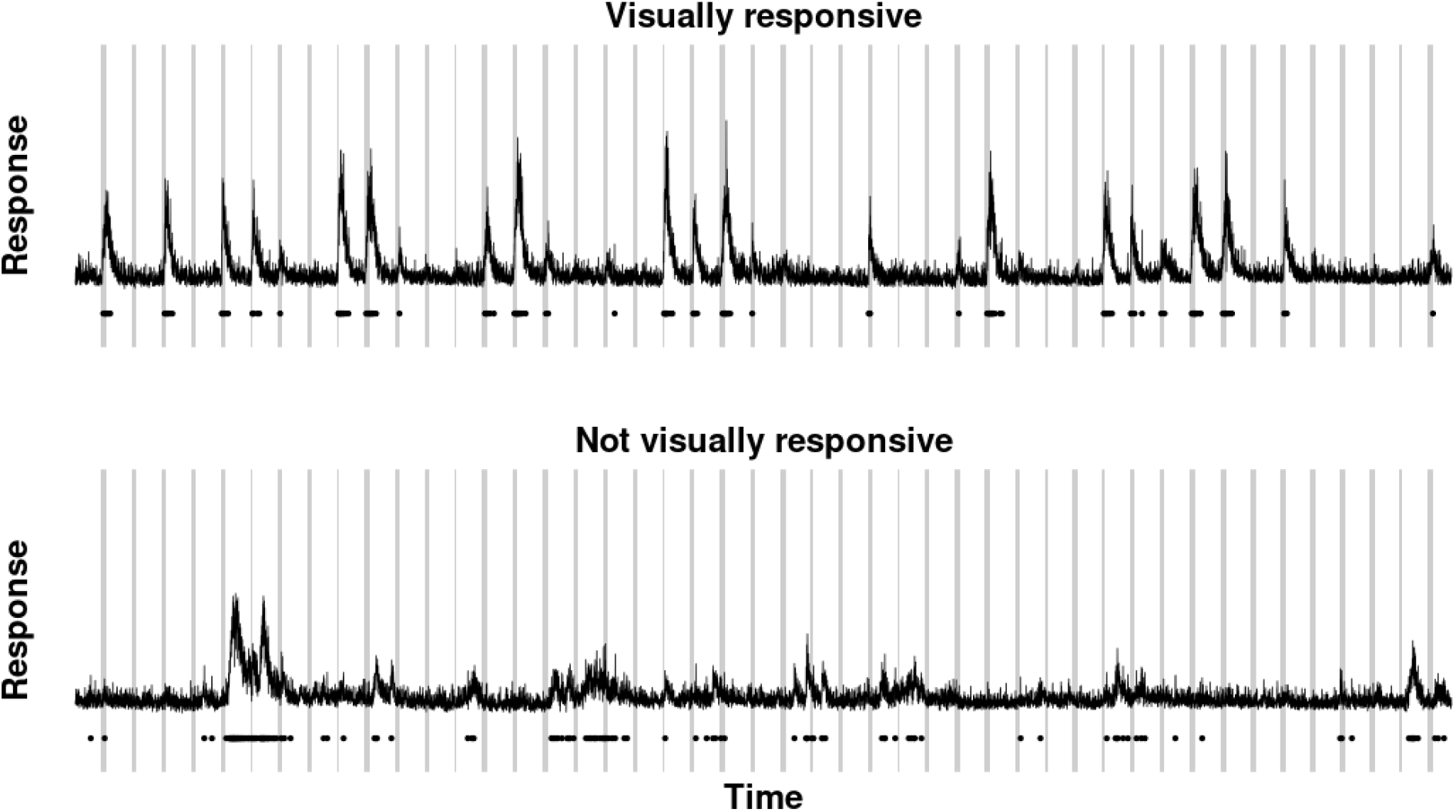
Example calcium transients from two cells during part of an imaging session. Grey bars indicate when a stimulus was being shown. Top panel shows a neuron classified as visually responsive, the bottom panel shows a neuron classified as not visually responsive. The dots under the calcium traces show time points when the neuron is inferred to be active (methods)

**Figure S2:**
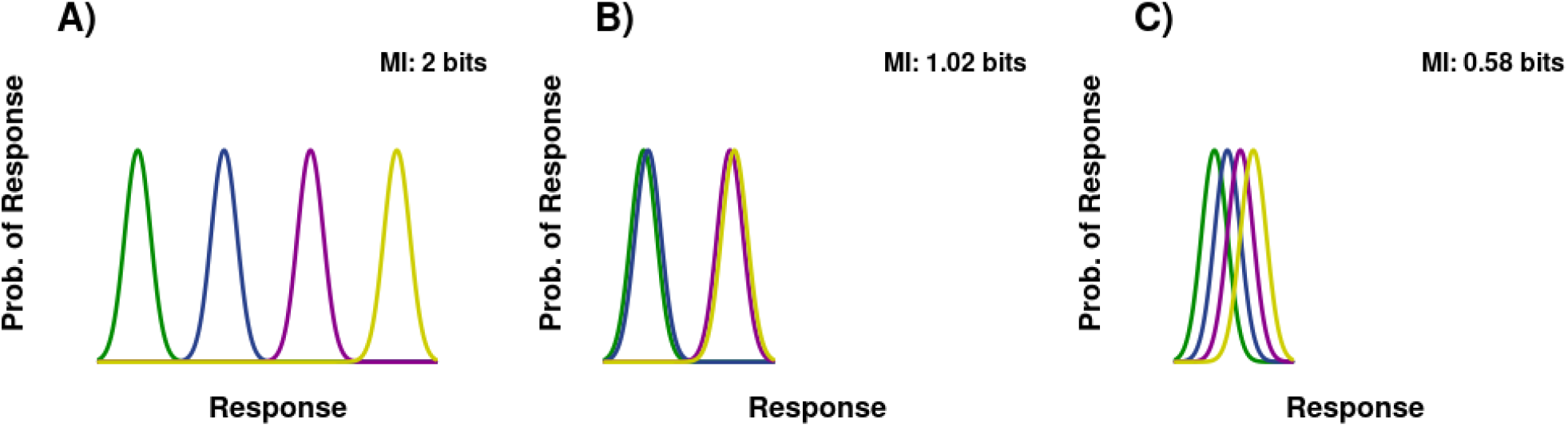
The number of bits a neuron encodes relates to how well separated the response distributions are. (**A**) The response distributions of a hypothetical neuron which encodes the maximum amount of information (2 bits). In this case all four response distributions are not overlapping, allowing each of the 4 stimuli to be decoded from a single neuron. (**B**) If response distributions overlap then the number of bits encoded decreases. In this case, the neuron responds in a very similar manner to the blue and green stimuli meaning the neuron cannot discriminate between them. The same is true for the yellow and purple stimuli. (**C**) If the response distributions are highly overlapping then only a small number of bits are encoded.

**Figure S3:**
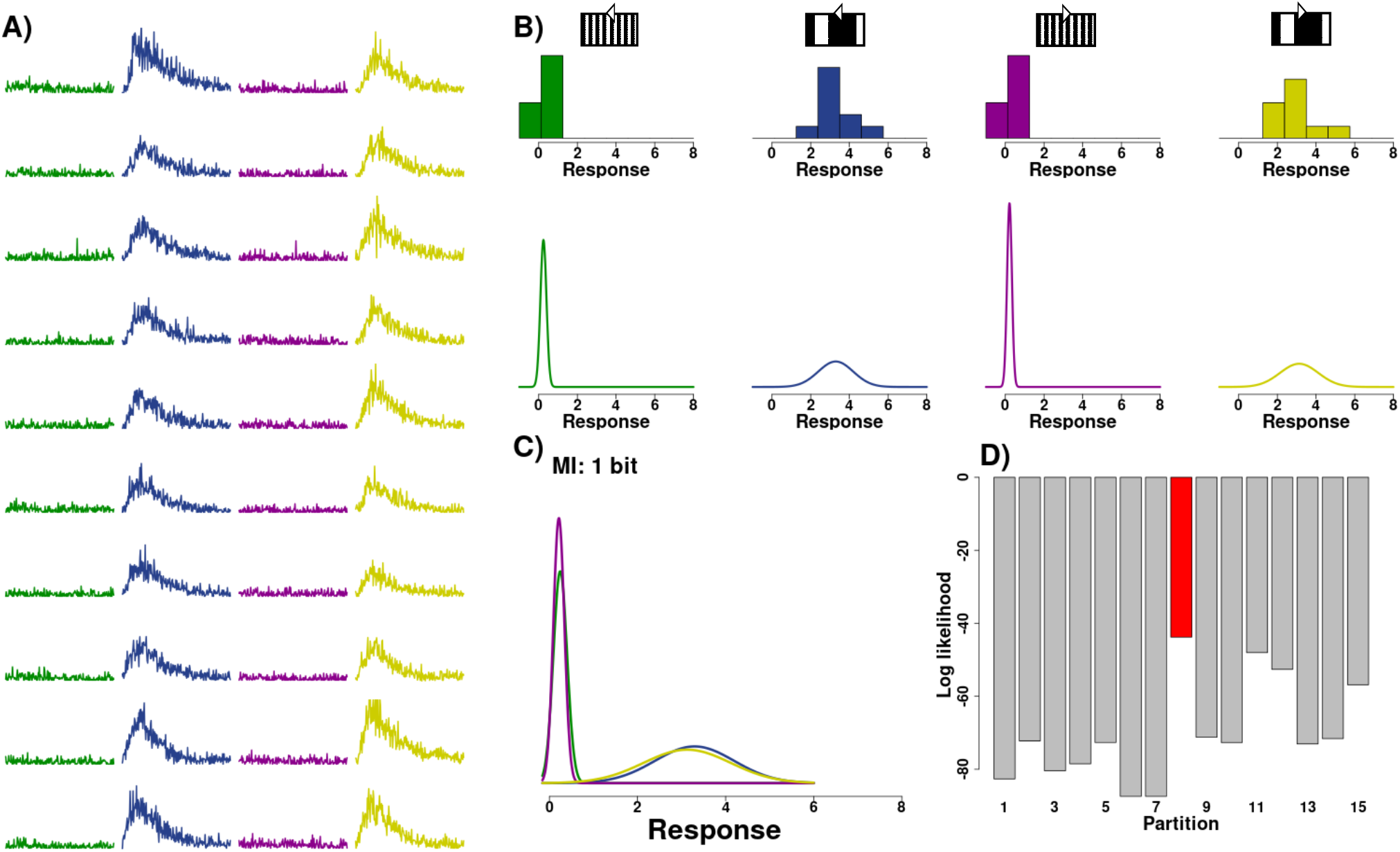
Workflow for calculating the mutual information of an example neuron. (**A**) Calcium transients of one example neuron to repetitions of the set of four grating stimuli, colour coded according to stimulus. (**B**) Above: histogram of the mean calcium fluorescence across each repetition for the four stimuli. Below: the estimated *P* (*R*|*S*) for each stimulus. (**C**) The *P* (*R*|*S*) for each stimulus overlaid with each other and the amount of mutual information the neuron was calculated to encode (MI: 1 bit). (**D**) The log likelihood for the neuron to belong to each partition of the set of stimuli (Table 1). The bar in red shows the partition with the highest log likelihood value, to which the neuron is then associated.

**Figure S4:**
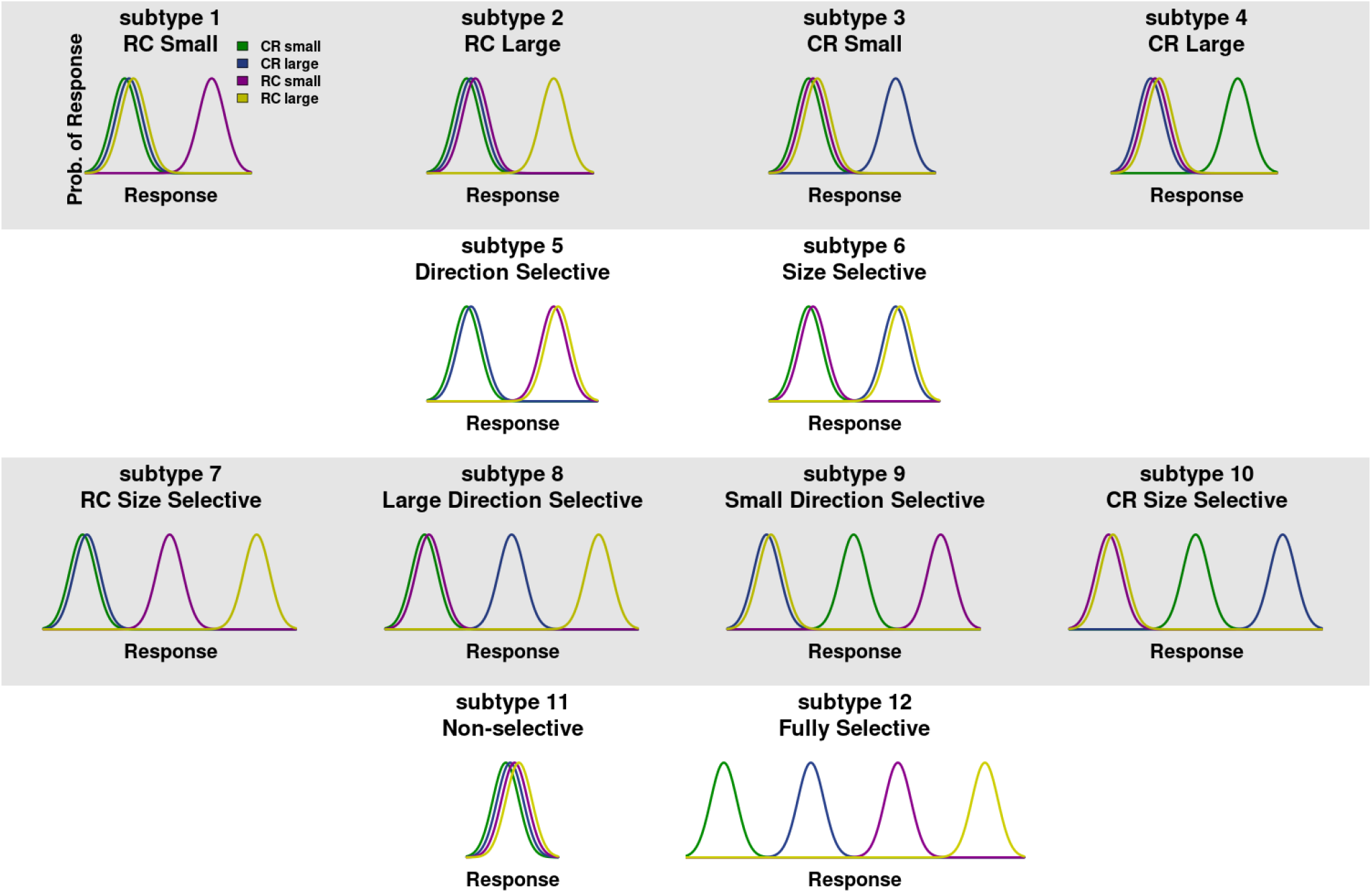
Schematic of functional neuronal subtypes. Neuronal subtypes are defined based on which stimuli, or types of stimuli, a neuron can best discriminate between. The figure shows schematics of idealised neuronal response distributions to the stimuli for each of the neuronal subtypes. CR: caudal-to-rostral direction of motion, RC: rostral-to-caudal direction of motion.

**Figure S5:**
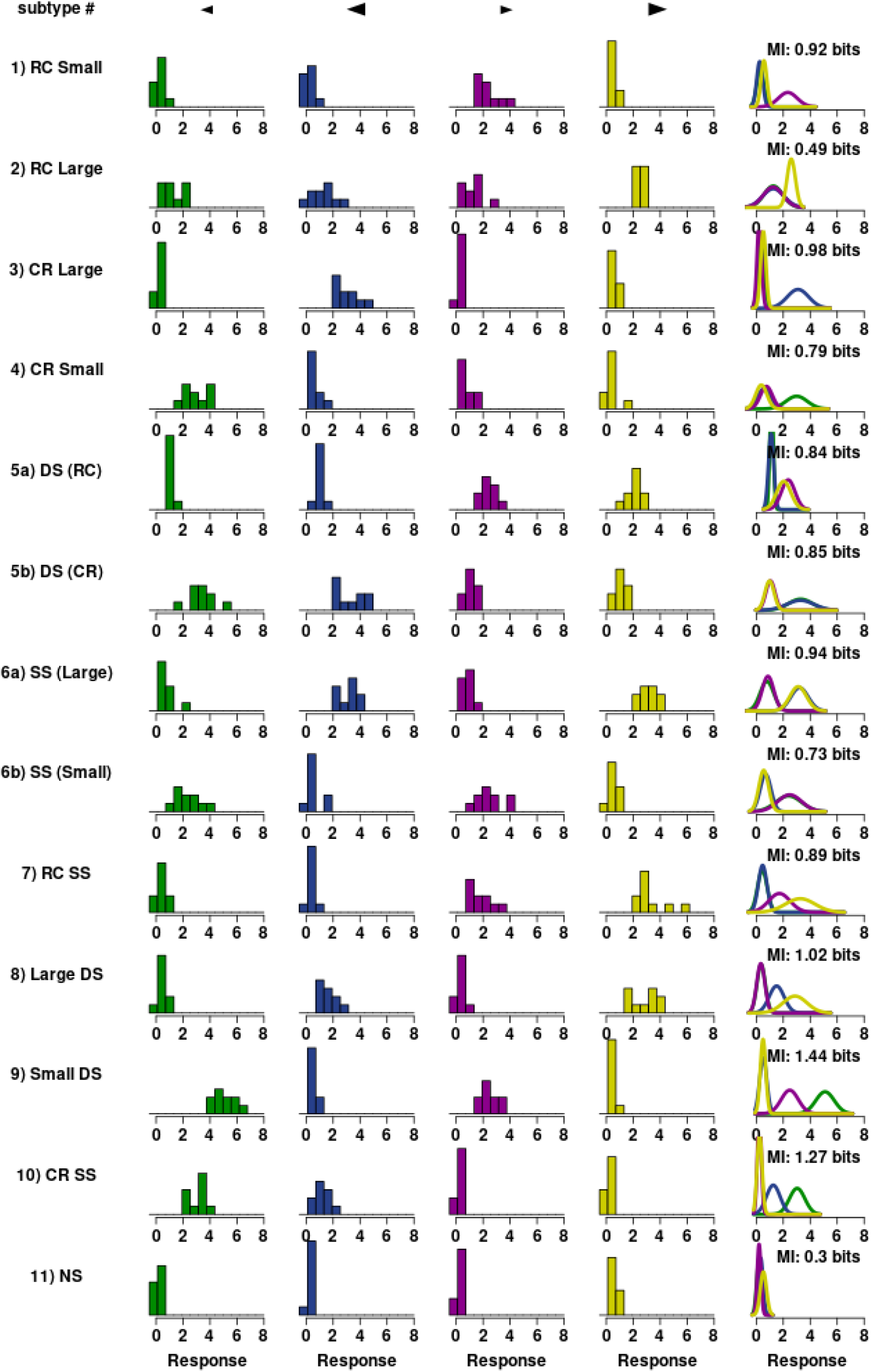
Example neurons from local motion subtypes. Each row shows an example local motion neuron that has been assigned to one of the subtypes. The histograms are the responses of that neuron to each stimulus, colour coded according to stimulus. The last column shows the overlaid estimated probability densities for the stimuli and how much information each neuron encodes. MI: mutual information, CR: caudal-to-rostral direction of motion, RC: rostral-to-caudal direction of motion, DS: direction-selective, SS: size-selective.

**Figure S6:**
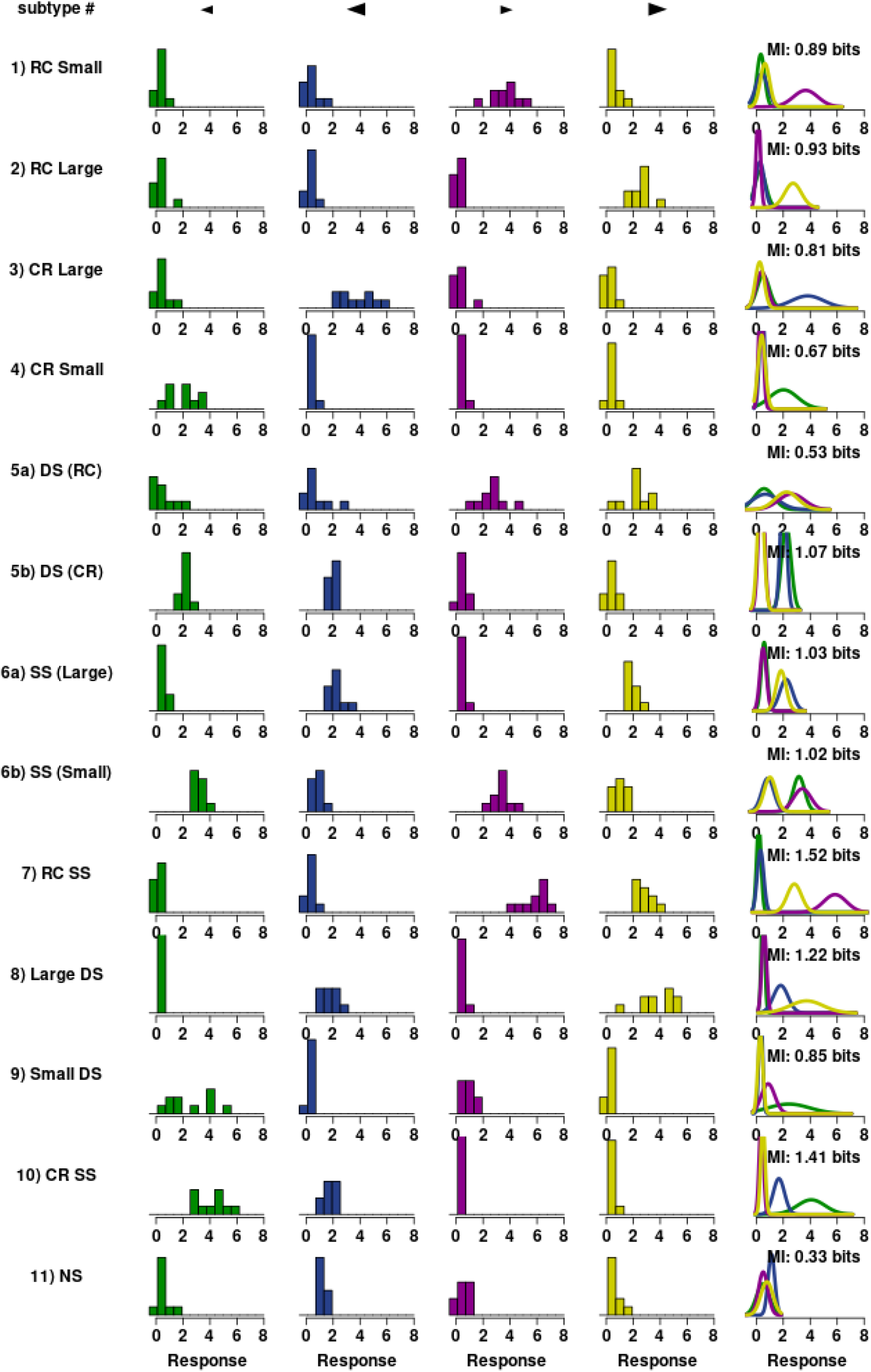
Example neurons from global motion subtypes. Each row shows an example global motion neuron that has been assigned to one of the subtypes. The histograms are the responses of that neuron to each stimulus, colour coded according to stimulus. The last column shows the overlaid estimated probability densities for the stimuli and how much information each neuron encodes. MI: mutual information, CR: caudal-to-rostral direction of motion, RC: rostral-to-caudal direction of motion, DS: direction-selective, SS: size-selective.

**Figure S7:**
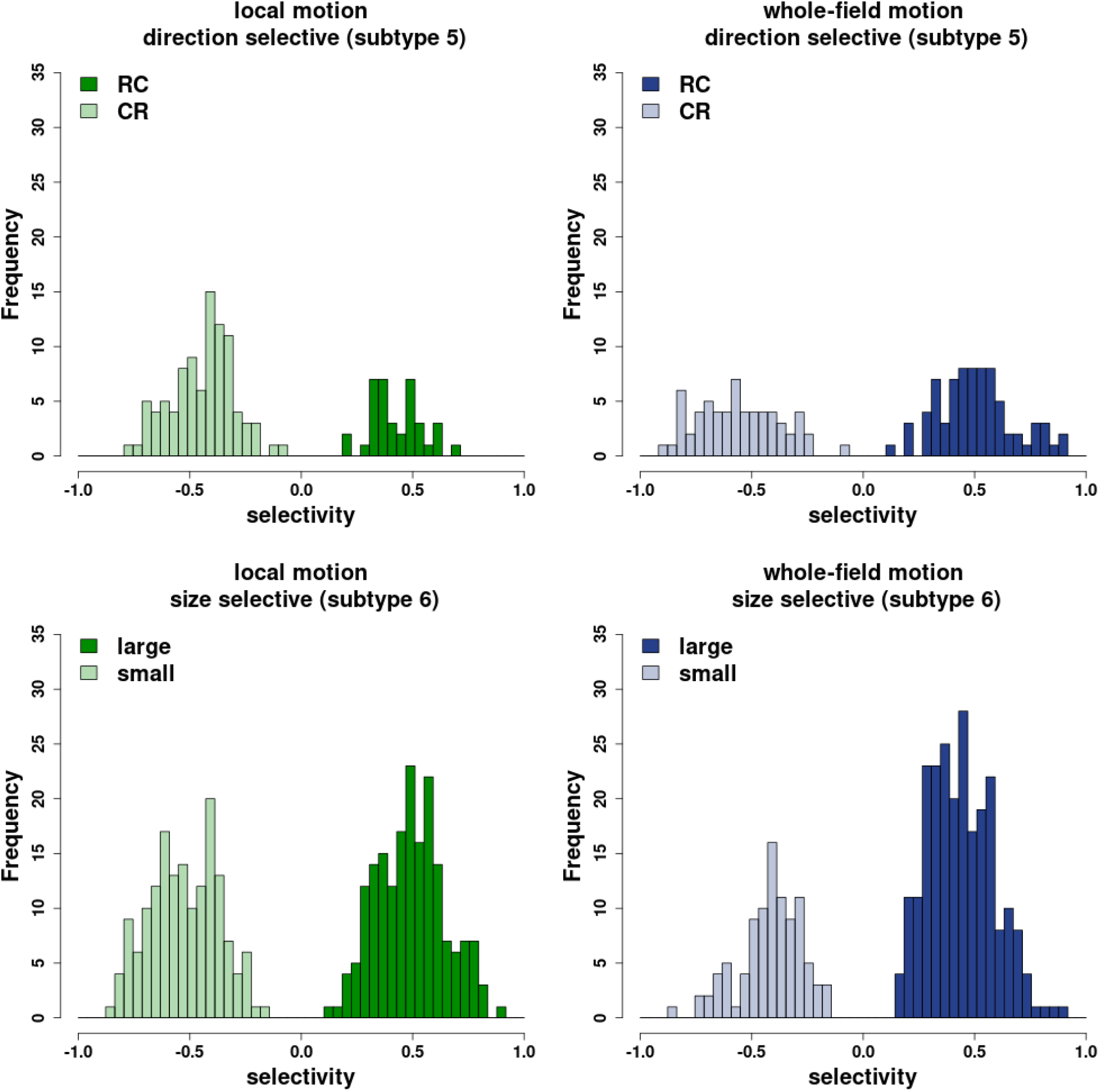
Direction and size selectivity of neurons. The top row shows the distribution of direction selectivity for neurons assigned to subtype 6, for either local motion (green) or whole-field motion (blue) encoding neurons. Neurons with a selectivity *>* 0 demonstrate a preference for rostral-caudal (RC) motion and were assigned to subtype 6a, whilst neurons with a selectivity *<* 0 demonstrate a preference for caudal-rostral (CR) motion and were assigned to subtype 6b. The bottom row shows the distribution of size selectivity for neurons assigned to subtype 7. Neurons with selectivity *>* 0 demonstrated a preference for large size stimuli (26) and were assigned to subtype 7a, whilst neurons with a selectivity *<* 0 demonstrate a preference for small size stimuli (5) and were assigned to subtype 7b.

**Figure S8:**
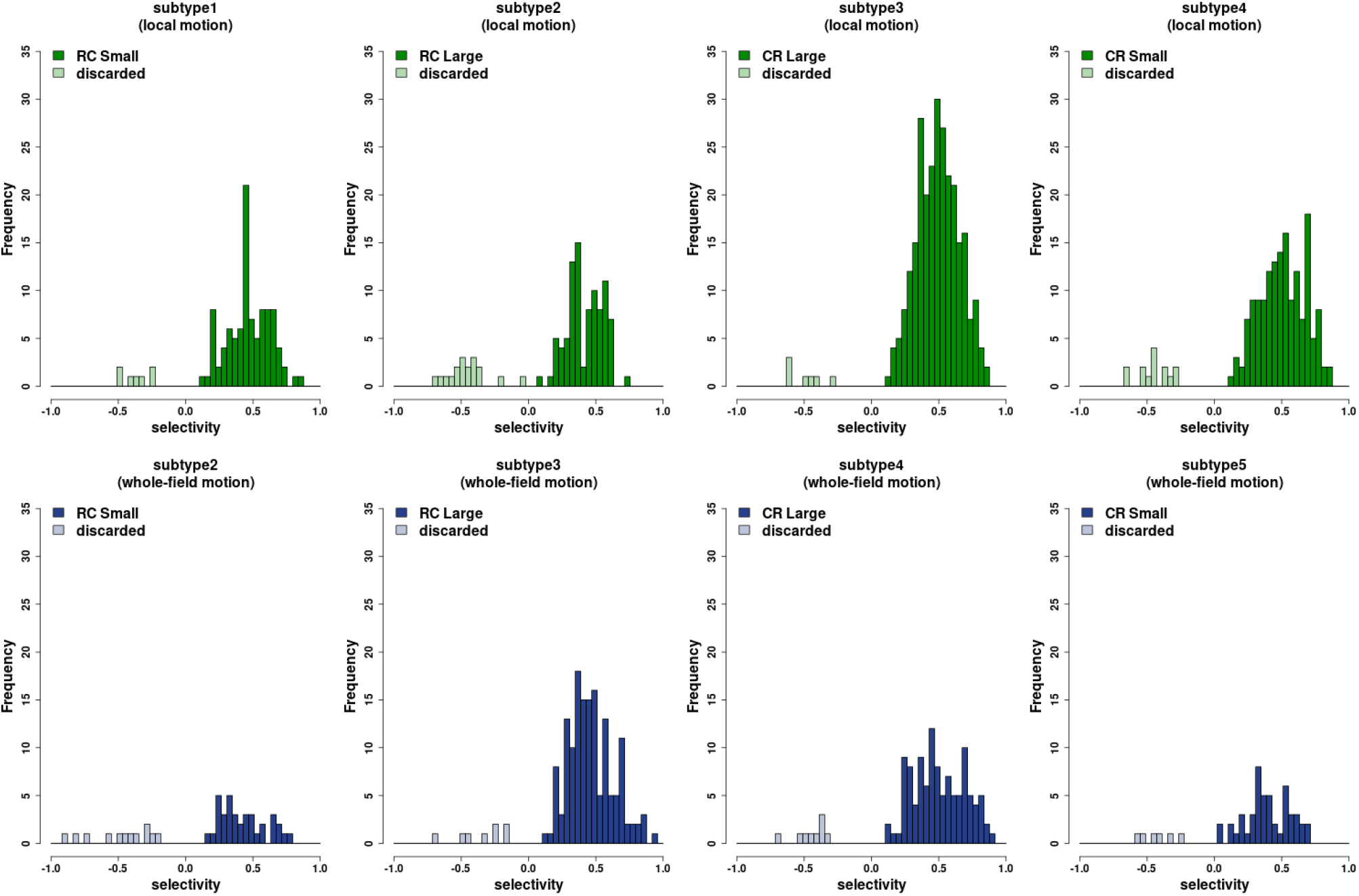
Single stimulus selectivity. Each plot shows the selectivity of neurons in subtypes 2-5 for stimuli in subset 2 vs subset 1 (Table 1). The top row shows local motion encoding neurons, the bottom row shows whole-field motion encoding neurons. Neurons with selectivity *>* 0 demonstrate a preference for the stimulus in subset 2. Neurons with selectivity *<* 0 demonstrate a preference for stimulus in subset 1, these neurons were discarded from further analysis.

